# Single-cell RNA sequencing reveals dysregulated fibroblast subclusters in prurigo nodularis

**DOI:** 10.1101/2023.01.29.526050

**Authors:** Jay R. Patel, Marina Z. Joel, Kevin K. Lee, Anusha Kambala, Hannah Cornman, Olusola Oladipo, Matthew Taylor, June Deng, Varsha Parthasarathy, Karen Cravero, Melika Marani, Ryan Zhao, Sreenidhi Sankararam, Ruixiang Li, Thomas Pritchard, Vito Rebecca, Madan M. Kwatra, Won Jin Ho, Xinzhong Dong, Sewon Kang, Shawn G. Kwatra

## Abstract

Prurigo nodularis (PN) is an intensely pruritic, chronic inflammatory skin disease that disproportionately affects black patients. However, the pathogenesis of PN is poorly understood. We performed single-cell transcriptomic profiling, ligand receptor analysis and cell trajectory analysis of 28,695 lesional and non-lesional PN skin cells to uncover disease-identifying cell compositions and genetic characteristics. We uncovered a dysregulated role for fibroblasts (FBs) and myofibroblasts as a key pathogenic element in PN, which were significantly increased in PN lesional skin. We defined seven unique subclusters of FBs in PN skin and observed a shift of PN lesional FBs towards a cancer-associated fibroblast (CAF)-like phenotype, with WNT5A+ CAFs increased in the skin of PN patients and similarly so in squamous cell carcinoma (SCC). A multi-center PN cohort study subsequently revealed an increased risk of SCC as well as additional CAF-associated malignancies in PN patients, including breast and colorectal cancers. Systemic fibroproliferative diseases were also upregulated in PN patients, including renal sclerosis and idiopathic pulmonary fibrosis. Ligand receptor analyses demonstrated increased FB1-derived WNT5A and periostin interactions with neuronal receptors MCAM and ITGAV, suggesting a fibroblast-neuronal axis in PN. Type I IFN responses in immune cells and increased angiogenesis/permeability in endothelial cells were also observed. As compared to atopic dermatitis (AD) and psoriasis (PSO) patients, increased mesenchymal dysregulation is unique to PN with an intermediate Th2/Th17 phenotype between atopic dermatitis and psoriasis. These findings identify a pathogenic role for CAFs in PN, including a novel targetable WNT5A+ fibroblast subpopulation and CAF-associated malignancies in PN patients.

## INTRODUCTION

Prurigo nodularis (PN) is a chronic inflammatory skin disease that is characterized by intensely pruritic nodules distributed on the trunk and extremities^1^. PN is associated with a reduction in overall quality of life and represents a significant health disparity^2,3^. Black patients are disproportionately affected with PN, and often suffer from a more severe disease presentation and are more likely to be hospitalized for complications^4–7^. Despite this significant burden, PN patients are often time resistant to treatment with conventional management^8,9^.

Previous studies have highlighted the role of cutaneous and systemic immune dysregulation by demonstrating involvement of the T-helper (Th)2-Th17-Th22 axis in the skin and an increase in interleukin (IL)-22 secreted by circulating blood CD4 and CD8 T cells^10,11^. CD4^+^, CD8^+^, γδ, and natural killer T-cells of these axes are increased in the circulation of PN patients^10,11^. Components of type 2 inflammation, IL-13 and periostin, are also elevated in PN blood^12^. Furthermore, neural dysfunction exists as PN lesional skin is characterized by increased dermal innervation and decreased epidermal nerve fiber density^13–15^. Recently, in a comparative bulk RNA-sequencing study, fibroproliferative gene activity was suggested as a unique pathogenic element of PN as compared to atopic dermatitis (AD) and psoriasis, two intensely pruritic inflammatory skin diseases^16^. However, PN remains poorly understood and is yet to be characterized on the single-cell level.

We thus performed single-cell transcriptomic profiling (scRNA-seq), an unbiased and indepth technique, to provide a detailed snapshot of the molecular landscape of PN. scRNA-seq was performed from lesional and non-lesional skin of PN Patients, with subsequent trajectory analysis, lineage tracing, and single-cell comparative transcriptomic mapping to inflammatory and neoplastic skin conditions. The findings were subsequently validated in multi-center cohort data to provide a robust assessment of PN pathogenesis.

This study discovers the presence of a unique fibroblast cell population in the skin of PN patients and uncovers the presence of a cancer-associated fibroblasts(CAF)-like phenotype in PN fibroblasts. This is supported by multi-center analyses demonstrating an increased risk of CAF-associated cancers in PN patients. Using ligand receptor analyses, we also described upregulated expression and signaling of CAF-derived WNT5A and periostin with neuronal expression of respective receptors MCAM and ITGAV. These findings demonstrate therapeutic discovery of treatment targets for the management of PN and provide a significant advance in our understanding of the pathogenesis of PN.

## RESULTS

### Identifying the single cell RNA-seq landscape of lesional and non-lesional PN skin

To investigate the single cell landscape of PN, we recruited 6 patients (5 African American and 1 Caucasian) with a clinical diagnosis of PN by a board-certified dermatologist and characteristic pruritic skin nodules (Fig. 1a). This study utilized lesional and non-lesional skin biopsies that were dissociated into single cell suspensions, barcoded, and processed using the Chromium 10x system. Data analysis was subsequently performed using single cell transcripts and compared with existing databases for relevant skin conditions (Fig. 1b). After quality control, a total of 28,695 cells were utilized for analysis. Cells were projected onto an unsupervised uniform manifold approximation and projection (UMAP) to visualize contributions by biopsy type and sample identity (Fig. 1c). Clustering was performed using the Louvain algorithm in Seurat with initial low resolution to identify major cellular clusters (Fig. 1d). Heatmaps for cellular clusters 0-21 were constructed for the top 10 differentially expressed genes (DEG) in each cluster to label and combine cell types (Fig. 1e). We identified major populations of B cells, mast cells, neurons, keratinocytes (KC), lymphatic endothelium (LEND), myeloid cells, natural killer T cells (NKT), endothelium (END), smooth muscle (SMC), FBs, and pericytes (PC), with the expression of representative markers displayed in Fig. 1f-g. B cells were defined based on the expression of CD79, IGHG1, and IGKC; mast cells expressed TPSAB1, TPSB2, and CTSG; neurons expressed NRXN1, S100B, GFRA3, and PLP1; KCs was subdivided into KC1 with higher keratins 4,14, and 15 while KC2 expressed KRT19 and DCD; LEND expressed LYVE1, PROX1, and MMRN1; myeloid cells expressed LYZ, HLA-DRA, CD68, and ITGAX; NKT cells expressed CD3D, CD4, CD8A, NKG7, XCL1, and KLRD1; END expressed VWF, PECAM1, and PLVAP; SMC expressed TPM2, MYL9, and MCAM; FBs expressed COL1A2, COL3A1, and DCN; PCs expressed RGS5, ACTA2, and NOTCH3 (Fig. 1f-g). The percentages of each major cellular cluster revealed trends for increased END and PC components (Fig. 1h). However, given the overlap of markers in some major clusters, we further refined the clusters with higher resolution unsupervised clustering, when possible, based on distinct expression profiles to avoid arbitrary clustering. The following populations were further subclustered: FB, PC, NKT, and myeloid. The FB population was split into seven (FB1-7) uniquely defined FB populations based on differentially expressed gene signatures (DEGs). The PC cluster was split into pericytes (PC1 and PC2) and myofibroblasts (mFBs), given the overlap of smooth muscle and collagen genes in the initial major cluster. NKT cells were split into T cell populations CD4 T cells, CD8 T cells, CD4 tissue-resident memory cells (CD4Trm), NK T cells (NKT), and unspecified T cells (Tun). Myeloid cells were split into three populations of conventional dendritic cells, all expressing CLEC10A (cDC2A-C), and two macrophage populations (Mac1-2). The percentages of these subclusters allowed for the identification of more refined differences in lesional and non-lesional PN skin (Fig. 1i). Correlation of major cellular clusters revealed positive correlations between PC/SMC, FB/neurons, and immune cell populations (NKT/B/myeloid/mast cells), (Fig. 1j). Correlation of subclusters accentuated immune cell populations, PC1-2/SMC/mFB, and FB1-7/neurons/KC1-2, (Fig. 1j).

**Fig. 1.**
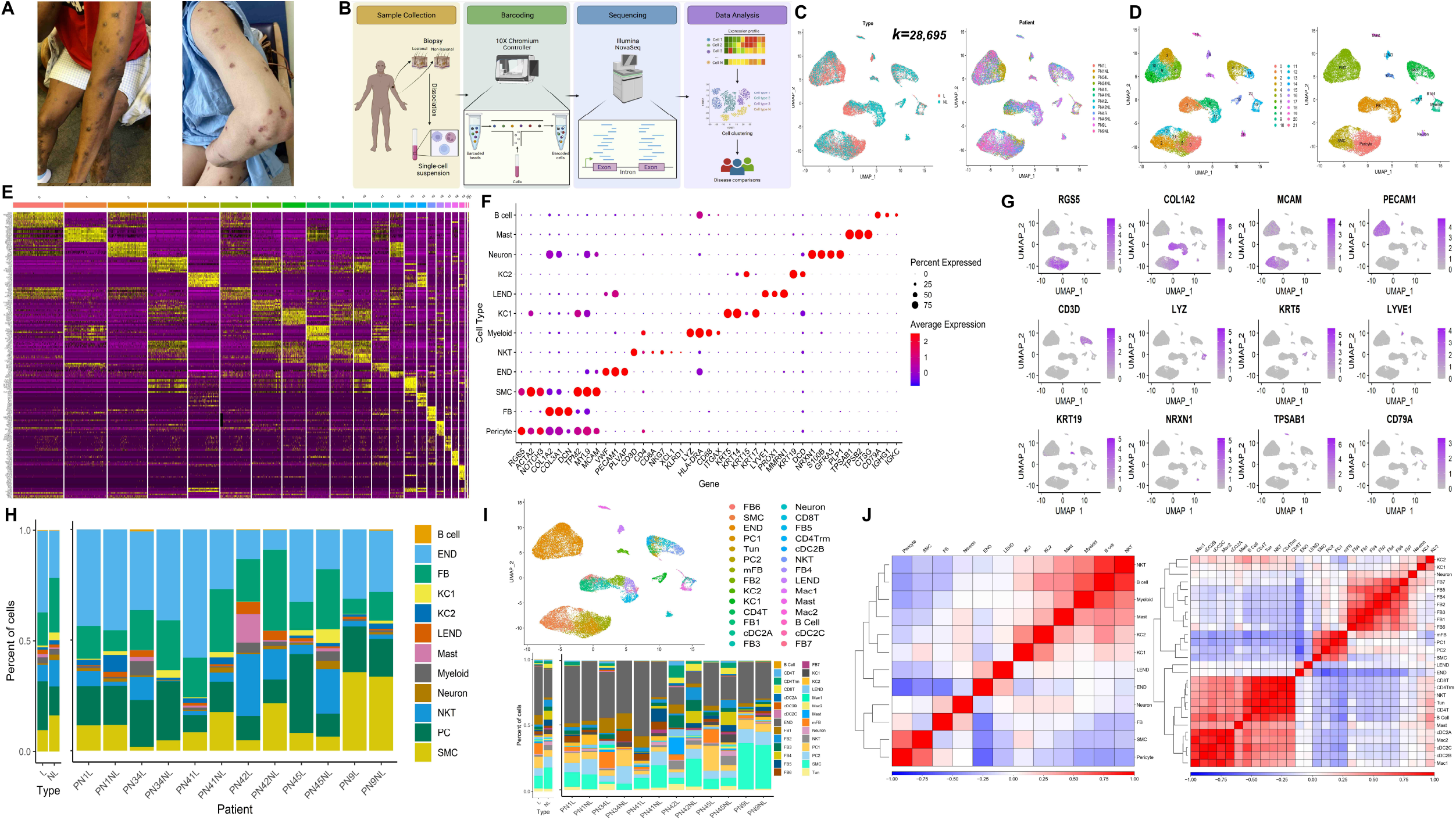
Single cell RNA landscape of Prurigo Nodularis. **a,** Representative patient photographs showing prurigo nodularis lesional (L) and adjacent non-lesional (NL) skin biopsied for downstream processing. **b,** Overview of scRNAseq study design demonstrating sample collection of PN L and NL skin followed by barcoding, RNA sequencing, and data analysis of single cell transcriptomics to identify cell types. Further characterization of PN performed using disease comparisons of existing scRNAseq databases. **c,** UMAP of PN cells by L and NL Type on the left with cells organized by sample on the right. **d,** Clusters 0-21 identified using the Louvain algorithm on the left with annotated clusters on the right. **e,** Heatmap of clusters 0-21 identifying top differentially expressed genes in each cluster. **f,** Dotplot of representative genes for each cell type cluster annotated. **g,** UMAP feature plot of representative genes corresponding to cell type annotation. **h,** Stacked barplot of the proportion of each cell type in each sample. **i,** UMAP of annotated subclusters identified on top with stacked barplot showing proportions of subclusters in each sample. **j,** Hierarchically clustered correlation matrices of cell types across all samples with original clusters on the left and subclusters on the right.

### Differential single cell characteristics of lesional versus non-lesional PN skin

We compared lesional vs. non-lesional PN skin to evaluate pathological changes contributing to lesion formation and itch. UMAP of major and subclusters showed differences in the FB, endothelial, and pericyte projections (Fig. 2a). Using pseudobulk RNAseq analysis, lesional skin was hierarchically separated from non-lesional skin (Fig. 2b). The top 20 most DEGs identified included genes related to fibroblast function and proliferation: COL4A2, FCN3, MMP1, PPRSS23, and WNT5A (Fig. 2b). Cluster-based compositional analysis was performed comparing loading coefficients of cell types showing increases in the mFB and cDC2A populations in lesional skin followed by an increase in CD4 T cells, END, and FB1 cluster (Fig. 2c). Traditional box-plot comparison also revealed similar trends (Fig. S1a). Hierarchical representation of compositional changes similarly showed a shift towards FB, T cell, cDC2, and endothelial populations (Fig. 2d). Cluster free compositional changes using the subtraction and Wilcox methods revealed similar trends, highlighting an increase in FB1, mFB, and END (Fig. 2e). Expression shifts were characterized by using normalized expression distance for each cell cluster; using this method mast cells and FB1 showed differential expression (Fig. 2f). Gene ontology analysis of each individual cluster highlighted differentially expressed pathways in lesional and non-lesional skin (Fig. 2g, S1b-c). There was an increase in morphogenesis, cell migration, proliferation, and collagen synthesis among FB subsets, SMC, END, and PC2. A type I interferon response was evident in T cells and myeloid cells. There were decreases in cellular detoxification and metabolic function in the END with Mac2 demonstrating decreased immune function. Individual genes are highlighted in an array of volcano plots (Fig. S1d). FB populations showed increased collagen synthesis (COL6A3 and COL12A), mFB expressed more POSTN, sMC/PC1-2 expressed more collagen (COL1A1, COL1A2, COL3A, and COL5A2), and the END was decreased in HLA related genes.

**Fig. 2.**
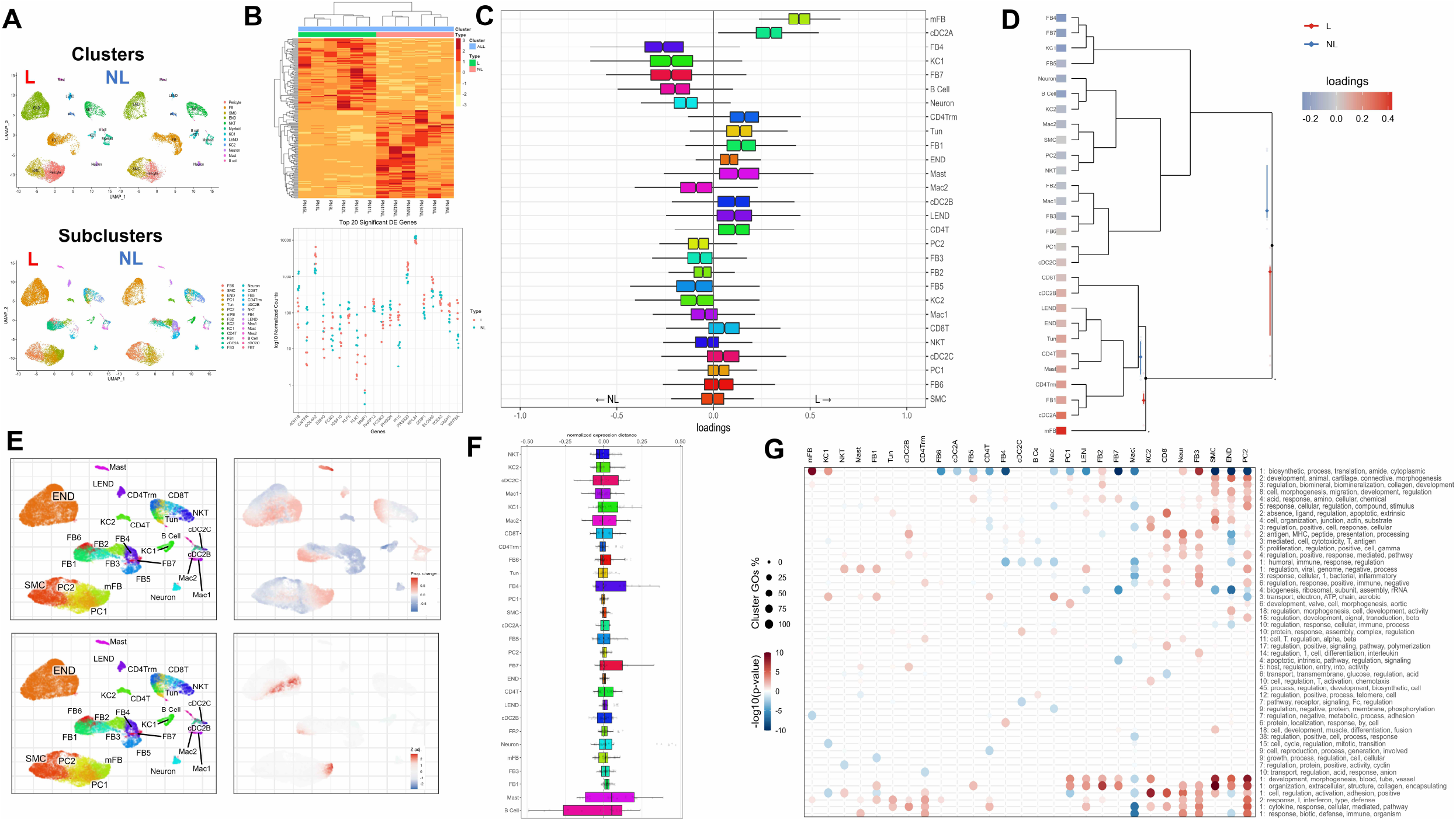
Differential single cell landscape of lesional vs non-lesional PN skin. **a,** UMAP plots of L and NL PN skin with major clusters on top and subclusters on the bottom. **b,** Pseudobulk RNAseq using single cell data identifies distinct L and NL transcriptomes as displayed by the heatmap. Top 20 differentially expressed genes between L and NL PN skin shown below. **c,** Compositional analysis of cell clusters showing differential cell loading coefficients of fibroblasts in L compared to NL skin. **d,** Hierarchical representation of compositional changes in L and NL skin. **e,** cluster free compositional changes based on subtraction (top) and wilcox testing (bottom, with significant z adjusted scores in red/blue). **f,** Expression differences calculated using normalized expression distance between L and NL skin for all cell clusters. **g,** Gene ontology heatmaps of top 50 pathways upregulated or downregulated in PN L vs NL skin.

### Lesional PN fibroblasts are shifted to cancer associated phenotype

Lesional skin demonstrated a shift toward the WNT5A+ FB1 population (Fig. 3a-b). This was further confirmed with immunofluorescence (IF), showing an increased colocalization of WNT5A and the fibroblast marker, vimentin, highlighting the FB1 subset in lesional PN skin biopsies (Figure 3c). Increased lesional SFRP2 and TNC was also noted in lesional and non-lesional fibroblasts compared to HC (Figure 3c). These FB populations were clustered based on DEGs that defined each subpopulation (Fig. 3d). Each subcluster was defined using representative markers as follows, FB1 (NKD2 and WNT5A), FB2: (CCL19, CD74, CLU), FB3: (SFRP2), FB4: (ITM2A), FB5: (SFRP1, TIMP3, ASPN), FB6: (MTRNR2L8), FB7: (APOD). Each L FB population, particularly FB1, was shifted toward a cancer associated fibroblast (CAF)-like phenotype, as defined by the genes (CTHRC1, NREP, WNT5A, POSTN, THY1, PMEPA1, CPXM1, TNC, GGT5) elevated across subclusters FB1-6 (Fig. 3e). An increased lesional type I interferon response was seen with genes (IFIG, IFI27, IFITM1, ISG15) across FB1. FB3 expressed increased IL32, and POSTN was increased throughout the lesional FBs. FB7, which was increased in non-lesional skin, expressed more CCL2 and SELENOP.

**Fig. 3.**
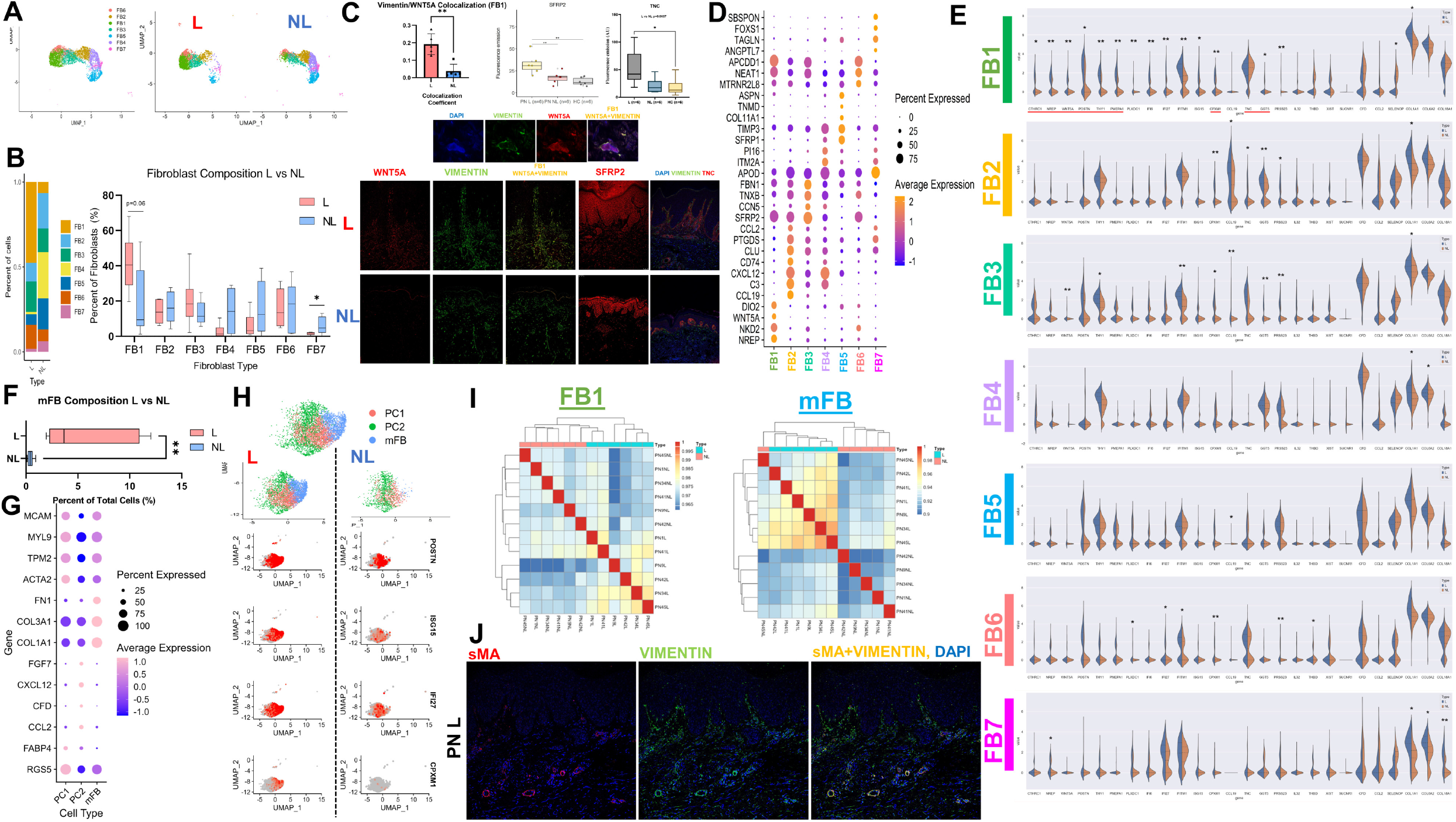
scRNAseq identifies cancer-associated fibroblast shifts in lesional compared to non-lesional PN skin. **a,** UMAP of fibroblasts FB1-FB7, separated by L and NL on the right. **b,** Stacked barplot showing differential composition of fibroblasts in L and NL PN (left). Statistical analysis via Mann Whitney test with * p<0.05. Barplot of L vs NL fibroblast subpopulations revealing trends for increased FB1 in L and increased FB7 in NL. **c,** Immunofluorescence imaging of L and NL PN skin revealing increased colocalization of WNT5A and Vimentin (FB1) cells using Manders colocalization coefficient in L PN and increased SFRP2/TNC expression in L and NL PN skin. Representative immunofluorescence images shown below. Statistical analysis via Mann Whitney test, ** p<0.01. **d,** Dotplot of representative genes identifying fibroblast subclusters. **e,** Split violin plots displaying differentially expressed genes in L vs NL fibroblast subclusters. All asterisks aside represent multiple t tests at a sample level using mean cellular expression, with *p<0.5 and ** p<0.01. **f,** Boxplot showing increased mFB composition in L compared to NL skin. Statistical analysis via Mann Whitney test with ** p<0.01. **g,** Dotplot of representative genes identifying subclusters (PC1, PC2, and mFB) split from the PC group. **h,** Feature Plots of PC1, PC2, and mFB revealing increased expression of genes POSTN, ISG15, IFI27, and CPXM1 in L cells compared to NL cells. **i,** Hierarchically clustered correlation matrix of FB1 (left) and mFB (right) revealing separation of L and NL skin. **j,** Immunofluorescence staining of L PN skin demonstrating overlap of sMA (red) and vimentin (green) with nuclei stained using DAPI (blue), highlighting high burden of mFB (yellow) in PN.

Given the expression profile of the PC major cluster, it was split into mFB and PCs. The mFB in lesional skin was greatly increased compared to non-lesional skin (Fig. 3f). PC subclusters PC1-2 and mFB were defined based on patterns of overlap of smooth muscle and collagen genes indicating myofibroblasts (Fig. 3g). PC1 was separated from PC2 based on the expression of chemokines CCL2, CXCL12, and FFG7. In lesional PN, PC1-2 and mFB populations expressed similar phenotypes to FB clusters in lesional skin, with increased POSTN, type I IFN (ISG15, IFI27), and CPXM1 (CAF gene), Figure 3h. Given the differences in FB1 and mFB, we attempted to segregate lesional and non-lesional skin with machine learning hierarchical clustering (Fig. 3i). Using this technique, FB1 perfectly separated lesional vs. non-lesional skin, whereas mFB misclassified 1 patient. Correlation analysis was performed on FB and PC clusters to identify populations influencing differentiation and proliferation of mFBs (Fig. S2a). In non-lesional skin, mFB positively correlated with PC1 (r=0.88, p<.05). Lesional skin was negatively correlated with FB4 (r=-0.94, p<0.05). Combining all samples resulted in a positive correlation with FB1 (r=0.71, p<0.05) and FB7 (r=0.62, p<0.05) but a negative correlation with FB4 (r=-0.73, p<0.05). Given the elevations in COL and POSTN, we investigated the differential expression of TGFB in mesenchymal cells and found increased trends throughout lesional populations of FB1, PC1-2, mFB, and SMC (Figure S2b).

### Differences in immune, endothelial, and neuronal lesional skin populations

The myeloid cluster was segregated into three cDC2 populations, cDC2A-C, and two macrophage populations, Mac1-2 (Fig. 4a-b). All cDC2 populations expressed CLEC10A with cDC2A defined by higher CD83, cDC2B with FCER1A, and cDC2C with FSCN1 and overall low expression of A/B markers. Mac1 aligned with more of an M1 phenotype expressing CD68, CD36, CCL13, and CCL18, while Mac2 was consistent with an M2 phenotype of IL10, CCL4, and CXCL2/3/8. L Mac2 expressed more chemokine CCL18, type I IFN genes, and TYMP (Fig. 4c). Along with increased count, cDC2A was also shifted toward a type I IFN phenotype in L skin (Fig. 4c). T cell populations were segregated as follows: NKT (NKG7), CD4Trm (CXCR6), CD8T (CD8), CD4T (CD4), and Tun was negative for distinguishing markers (Fig. 4d). The T cell repertoire in this study did not show significant shifts in lesional vs. non-lesional skin. However, there were trends for increased GNLY in CD8 T cells (p=0.23) and increased ISG15, TIGIT, and CTLA4 in CD4Trm (p=0.052, 0.053, 0.17; Fig. 4f). The endothelial populations END and LEND both expressed markers of angiogenesis and proliferation (Fig. 4g). These populations also expressed increased LGALS in lesional skin, related to immune migration and extravasation. The overall angiogenesis score was considerably increased in lesional compared to non-lesional skin (Fig. 4h). Cell Chat ligand receptor analysis on these populations revealed increased pro-angiogenesis signaling via ANGPT/ANGPTL and immune migration via GALECTIN (Fig. 4h). DEG analysis in neurons identified trends for increased TAFA5. In contrast, ligand receptor analysis revealed increased midkine (MK) signaling (Fig. 4i). Mast cells in lesional skin notably expressed identification markers CD9/CD52, chemotaxis markers LGALS3/LTC4S, and type I IFN markers IFITM1/2 (Fig. 4j).

**Fig. 4.**
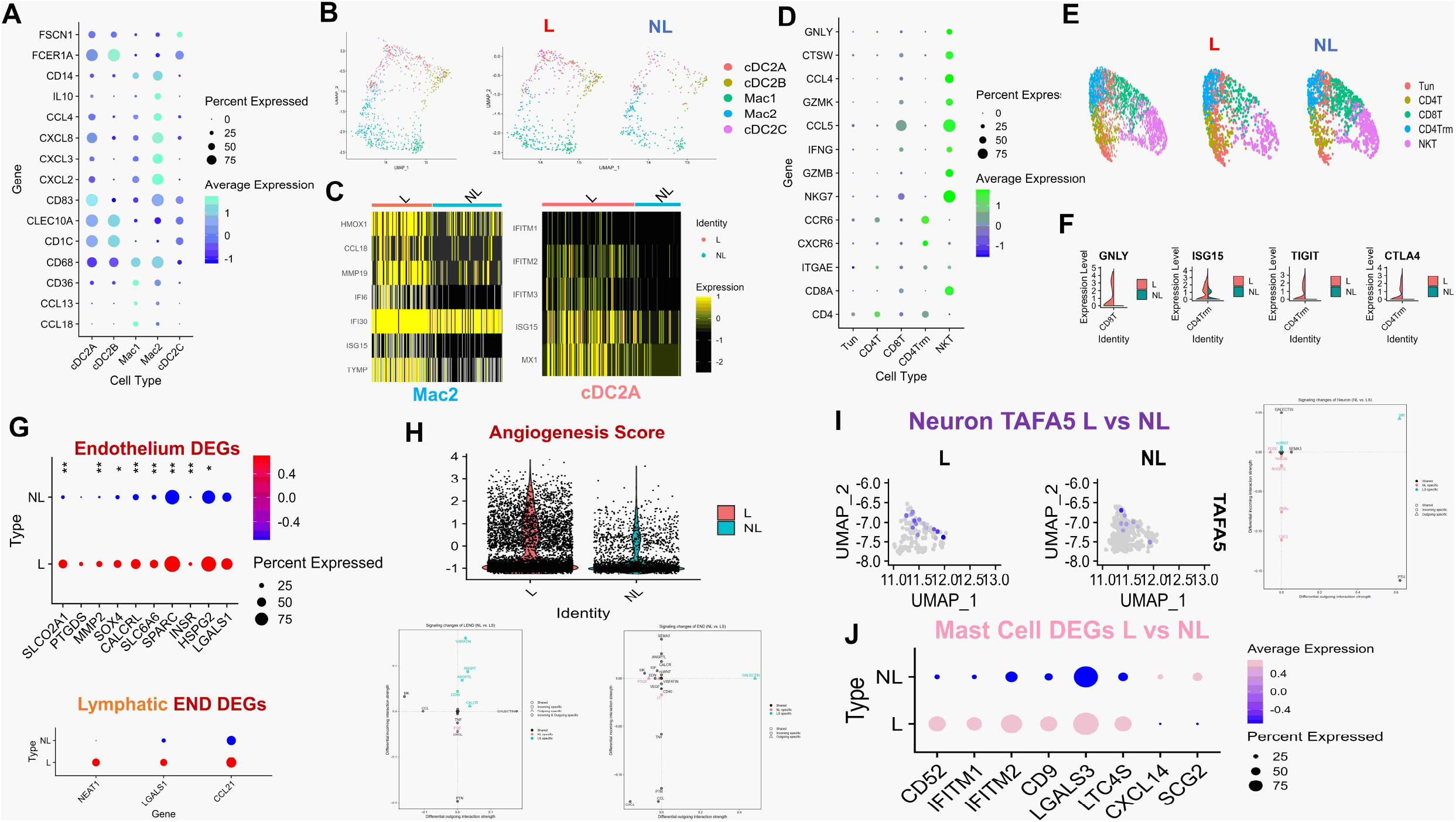
Single cell differences in lesional PN immune, endothelial, and neural populations. **a,** Dotplot of representative genes highlighting myeloid subclusters cDC2A, cDC2B, cDC2C, Mac1, and Mac2. **b,** UMAP of myeloid lineage (left), further split into L and NL (right). **c,** Heatmap of L vs NL Mac2 and cDC2A subclusters highlighting increased type I interferon genes in L populations. **d,** Dotplot of representative genes highlighting NKT cell subclusters of Tun, CD4T, CD8T, CD4Trm, and NKT. **e,** UMAP of NKT major cluster (left), further split into L and NL (right). **f,** Split violin plots displaying increased expression in L PN skin of GNLY in CD8T cells and ISG15/TIGIT/CTLA4 in CD4Trm. **g,** Dotplot of differentially expressed angiogenesis related genes in endothelium and lymphatic endothelium. **h,** Violin plots of angiogenesis score showing increased expression in L skin. **i,** Feature plot displaying increased TAFA5 expression in neural lineage cells in L skin. **j,** Dotplot of DEGs in L Mast cells. All comparisons between NL and L gene expression at a cellular level represent p<0.05 using wilcoxon rank sum test. All asterisks represent multiple t tests at a sample level using mean cellular expression, with *p<0.5 and ** p<0.01.

### Ligand receptor analysis of PN

We implemented ligand receptor analysis better to understand the transcriptome’s functional significance in PN skin. Lesional and non-lesional ligand-receptor analysis was performed using CellChat, with clear differences in the number and strength of interactions between lesional and non-lesional skin (Fig. S3a-d). FB populations interacted less and had a lower relative strength in non-lesional skin (Fig. S3b). The cDC2 subsets interacted more and had greater strengths of interactions in non-lesional skin (Fig. S3b). When comparing the overall information flow in lesional and non-lesional skin, key lesional skin pathways included periostin, ncWNT, VISFATIN, and TWEAK (Fig. 5a). Dominant non-lesional skin pathways included lymphotoxin, pleiotrophin, and complement (Fig. 5a). In lesional PN skin, overall signaling of periostin was increased in FB subsets. In contrast, overall signaling of visfatin was increased in FB4 and LEND (Fig. 5b). Non-lesional skin exhibited increased overall signaling of the lymphotoxin pathway in CD4Trm, pleiotropin in FB2/4/5, and complement in FB2-4 and cDC2 cells (Fig. 5b). Pathways were clustered using k means, with 4 clusters identified for functional and structural pathways (Fig. S3e). Cluster 1 in lesional skin included periostin associated with FGF and GAS. Functional cluster 2 included TNF for lesional skin, while structural cluster 2 included ncWNT for lesional skin associated with EDN and SEMA3. Functional cluster 4 included MIF associated with CXCL and MK in lesional skin. Cell patterns and communication patterns were identified using k means in lesional skin (Fig. 5c, S3f). Outgoing cell patterns included T cells/mast/B cells, Mac/cDC2 cells, and FB/END/neurons. Outgoing communication patterns in lesional skin included CD40/IFNII/LIGHT/FL3, VEGF/EGF/IL1/APRIL, ncWNT/complement/MK/PTN/CALCR, and TWEAK/KIT/NIT. Incoming cell patterns included END/LEND, FB/SMC/PC, T cells/B/Mast, and Mac/cDC2. Incoming communication patterns included CSF/complement/IL1/FILT3, ANGPT/ANGPTL/VEGF/CALCR, ncWNT/EDN/GRN/GAS/PROS, EGF/TWEAK, and APRIL/BAFF/CXCL. Single cell signaling changes occurred in FB populations with lesional skin having increased periostin and ncWNT signaling in FB1 while FB3/mFB had increased periostin signaling (Fig. S3g). Periostin signaling network in lesional skin revealed increased interactions between FB subpopulations, with FB1/3/5 being major mediators and influencers (Fig. 5d-e). Increased periostin throughout lesional PN compared to HC was confirmed via IF imaging (Figure 5e). Periostin interacted with ITGAV and ITGB5 in lesional skin with FB populations expressing periostin and the receptor ITGB5, while neurons were major expressors of the ligand ITGAV (Fig. 5f). Confirmatory IF imaging revealed neuronal B-III tubulin expressing ITGAV throughout lesional PN skin (Figure 5f). The ncWNT signaling network in lesional skin revealed a dominant role for FB1, while this was not evident in non-lesional skin (Fig. 5g). In lesional skin, WNT5A was associated with receptors MCAM, FZD1, FZD4, and FZD7, whereas in non-lesional skin WNT11 interacted with FZD1 and FZD4 (Fig. 5h-i). In lesional skin, FB1 predominantly expressed WNT5A and interacted with MCAM on multiple cell types, including neurons, SMC, PC, and mFBs (Fig. 5h). IF confirmed this expression in neurons with colocalization of B-III tubulin and MCAM in lesional PN skin (Figure 5j). FB1 was the major sender and influencer in lesional skin, with populations of neurons, PC, and mFB being receivers. Non-lesional skin differed in complement activation with FB expressing C3 and interacting with ITGAX/ITGAM/ITGB2 on cDC2 and T cell populations (Fig. S3h). FB2-4 were senders, and FB4 was a major influencer in the non-lesional complement pathway.

**Fig. 5.**
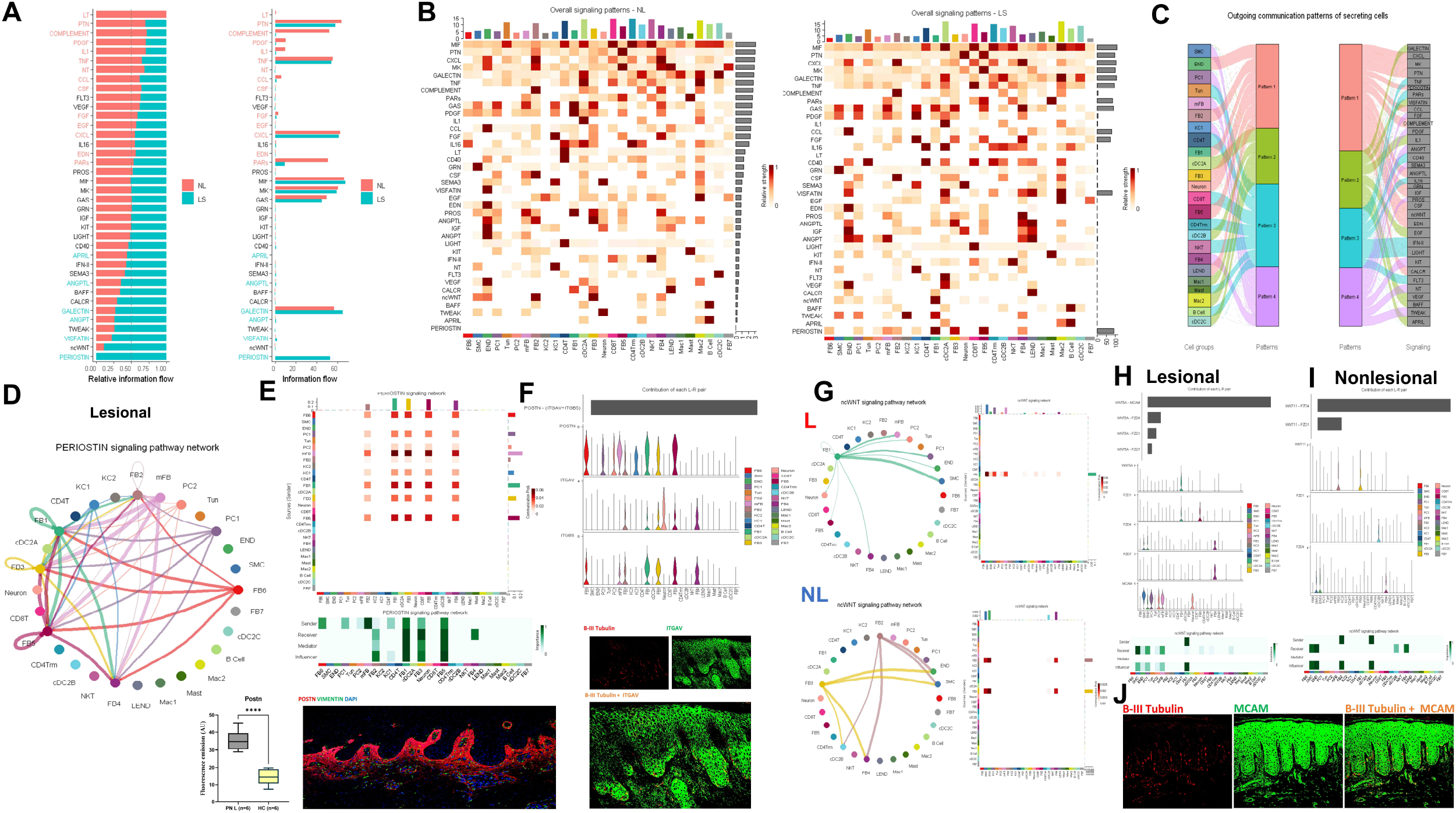
Ligand Receptor analysis of lesional and non lesional PN. **a,** Split barplots of information flow in L and NL PN revealing increased activity of periostin and wnt5a in L PN skin. **b,** Heatmaps of overall signaling patterns in NL (top) and L (bottom) PN **c,** Outgoing (secretion) pathway analysis of ligand receptors in L PN skin. Cell patterns and communication patterns defined using k means clustering. Flow charts display cell group and pathway contribution to cellular and pathway patterns. **d,** Circle plot of periostin pathway in L PN demonstrating FB activity. **e,** Heatmap of periostin pathway in L PN demonstrating FB activity (top). Heatmap of sender, receiver, mediator, and influencer categories demonstrating FB1 predominance in periostin signaling (middle). Confirmatory immunofluorescence staining of L PN demonstrating increased POSTN compared to HC (red) surrounding vimentin (green) with nuclei stained with DAPI (blue), (bottom). **f,** Violin plots of POSTN ligand and receptors ITGAV + ITGB5 showing expression of ligands in FB populations with ITGAV highest in neurons. IF showing colocalization of B-III tubulin (red) and ITGAV (green) in L PN skin (yellow) **g,** Circle plots and heatmaps of ncWNT pathway in L (top) and NL (bottom) PN, showing FB1 dominance in L skin. **h,** L characterization of ncWNT ligands and receptors MCAM, FZD1, FZD4, and FZD7 with highest contribution being WNT5A + MCAM in L skin (top). Violin plots of WNT5A and receptors showing high expression of ligands in FB1 with MCAM in PC, SMC, mFB, and Neurons (middle). Heatmap of sender, receiver, mediator, and influencer categories demonstrating for L WNT5A signaling (bottom). **i,** NL characterization of ncWNT ligands and receptors, FZD1 and FZD4 with highest contribution being WNT11 + FZD4 in NL skin (top). Violin plots of WNT11 and receptors (middle). Heatmap of sender, receiver, mediator, and influencer categories demonstrating for NL WNT11 signaling (bottom). **j,** IF imaging revealing colocalization of B-III tubulin (red) with MCAM(green) in PN L skin (yellow).

### RNA velocity and pseudotime analysis reveal separate FB differentiation patterns with distinct transcriptome progression in lesional subsets

To better define the relationship between cell populations in PN skin, we performed RNA velocity analysis, an approach that utilizes spliced and unspliced mRNA counts to predict the directionality and speed of cell state transitions. We projected RNA velocity onto UMAP and visualized velocity pseudotime derived from RNA velocity, which indicated that FBs were more terminal in differentiation than all other cell populations (Fig. 6a, b). RNA velocity-PAGA revealed epithelial to mesenchymal transition as END velocity vectors pointed toward the PC1 cluster and KC1 velocity vectors pointed toward the FB7 population (Fig. 6c). PAGA velocity analyses also showed a transition from PC1 to mFB, as well as separate FB differentiation trajectories: the first involving KC1, FB7, FB5, FB4, FB2, FB6, and the second involving mFB, FB1, FB3, both listed in approximate differentiation order. WNT5A and POSTN exhibited high RNA velocity and gene expression in FB, PC, and mFB clusters (Fig. S4a). Using CellRank, we determined seven unique cell lineages (END, FB, Mast, NKT, neuron, SMC, cDC), and computed a fate map for the FB lineage which demonstrated that KC, PC, mFB, and SMC cells also had high absorption probabilities for the FB lineage, indicating that those cells were likely to develop toward a FB terminal state (Fig. 6d). Top FB lineage driver genes were COL6A2, COL1A2, COL6A1, COL3A1 (*p* < 0.001) (Fig. 6e). Dynamic gene expression across unique cell lineages is demonstrated in Fig 6f, highlighting WNT5A and POSTN expression in FB with MCAM/ITGAV expression in neurons.

**Fig. 6.**
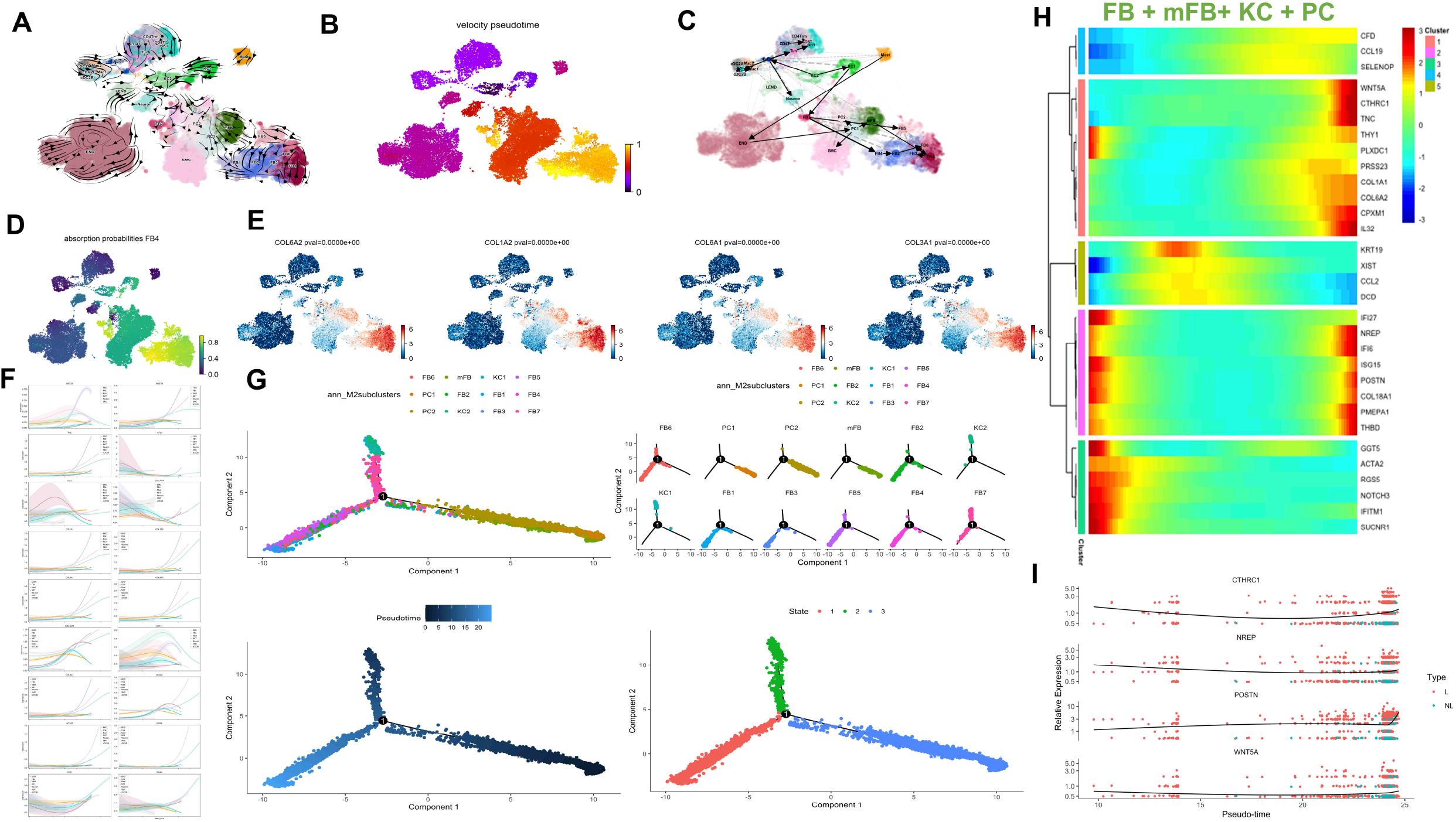
Trajectory analysis via RNA velocity and pseudotime reveals separate fibroblast differentiation patterns with distinct transcriptome progression in lesional subsets. **a,** RNA velocity fields projected onto UMAP. The streamlines indicate the directions of cell differentiation as inferred from RNA velocity estimations. **b,** Velocity pseudotime, derived from RNA velocity, is visualized in UMAP. **c,** RNA velocity-PAGA graph. Arrows representing direction of cells’ flow of PAGA-velocity were projected onto UMAP **d,** Computed fated map for the fibroblast lineage with absorption probabilities representing how likely each cell is to develop toward a fibroblast terminal state. Cell clusters included in the fibroblast lineage are PC, KC, SMC, and FB. **e,** Top lineage driver genes for the fibroblast lineage are plotted. **f,** Gene expression trends are plotted along separate lineages (END, FB, Mast, NKT, Neuron, SMC, cDC). Each lineage is defined via its lineage weights computed via CellRank. **g,** Pseudotime trajectory visualized in reduced dimensional space colored by cell type (top left), separated by cell type (top right), colored by pseudotime (bottom left), colored by cell state (bottom right). **h,** Heatmap of different blocks of DEGs along the pseudotime trajectory for FB, PC, mFB, KC. **i,** Plotting of gene expression in FB1 cells for select genes over pseudotime.

To further elucidate the pattern of dynamical cell state transitions, we performed pseudotime analyses for PC, mFB, and KC cells, populations from the FB lineage defined by RNA velocity analyses. Pseudotime trajectory visualization showed that cells occupied three major states and that State 3 (PC and mFB dominated) was the earliest differentiated, followed by State 2 (KC and FB-dominated), and State 1 (FB-dominated), the terminally differentiated state (Fig. 6g). These results align with earlier RNA velocity analyses indicating that FBs are most terminally differentiated and that KC1 cells differentiate towards FB7. Analyses of gene expression over pseudotime in FB1 cells showed distinct transcriptome progression in lesional vs. non-lesional subsets; WNT5A and CTHRC1 were present in both early and late differentiation throughout lesional FB1 but present only in late differentiation for non-lesional FB1 (Fig. S4b). Heatmap of gene expression over pseudotime for combined FB, PC, mFB, and KC cells demonstrated that PC genes (RGS5, ACTA2) were expressed earliest, followed by KC genes (KRT19, DCD), and finally FB genes (COL1A1, POSTN) (Fig. 6h). Across pseudotime, lesional skin expressed more CAF genes (CTHRC1, NREP, POSTN, and WNT5A, Fig 6i).

### A unique single cell signature of PN as compared to AD and PSO with a FB predominance

To validate the clustering of a single cell in PN samples and compare across different conditions and HCs, we utilized existing scRNAseq databases to obtain 6 HC, 4 AD, and 3 PSO biopsies. All samples were integrated using Seurat to reduce variations across studies. Using the louvian clustering algorithm, UMAP showed an alignment of conditions with differential cell populations (Fig. 7a, b). We performed a similar process of identifying major clusters followed by optimal sub clustering using adjusted resolution parameters. Major clusters (subclusters) included PC (PC1-2, mFB), END, SMC, FB(FB1-8), NKT (Tun, NKT, CD8T, Treg, CD4Trm, and proliferating CD4 T cells (CD4pro), myeloid (Mac1-3), LEND, KC, neuron, mast, and B cell. The FB populations that aligned with the PN cohort included FB1/2/3/5/7. Previously FB6 was integrated into FB1 with the combined dataset due to similarities in gene expression identified with the larger dataset. FB4 and FB8 were newly identified with the integrated dataset. Major cellular compositions differed, with PN having increased FB/END, AD having increased myeloid, Mast/B cells/NKT, and PSO having increased KC/NKT (Fig. 7c, d). Subclusters and UMAPs showed differences between conditions within each major cluster (Fig. 7e). The FB subsets FB1-8 showed increased FB1 in PN relative to AD and PSO and increased FB3 in AD relative to PN and PSO (Fig. 7f). Compared to HCs, PN showed an increase in mFB, PC1/2, FB1, and END (Fig. S5a-d). Top upregulated pathways included angiogenesis and collagen processing within mesenchymal and endothelial populations (Fig. S5g-h). Expressional differences existed in FB subsets, neurons, END, and SMC (Fig. S5e-f). To identify systemic disease, PN non-lesional skin was compared to HC skin, highlighting similar increases in mFB, PC1/2, SMC, FB1, and FB4 (Fig. S6a-d). Expressional shifts were predominant in FB1/3/4/7, SMC, and KCs (Fig. S6e). PN non-lesional skin had increases in PC metabolic pathways, CD4Trm antigen presentation and activation, and B cell response (Fig. S6f). FB1/3/7 all showed increased expression of genes RMRP, CCN2, STC1, PLCG2, NOVA1, CCL21, PTGDS, TNC, APOD, ID1, NR4A1, ID3, IGF2, CD74, FTH1, IGFBP3, EDNRA, POSTN, CRABP1, TFAP2A, and SFRP2 (Fig. S6g). Non-lesional neurons expressed increased markers of neural activity, and non-lesional CD4Trm’s expressed higher IL31 (Fig. S6h). Complete DEGs for each cell type are demonstrated in Fig. S6j). Comparison to inflammatory diseases AD and PSO identified a distinct signature in PN. Compared to AD, lesional PN skin had increased mFB, PC1, FB1, and neurons, while AD had increased immune cells, including macrophages, T cells, dendritic cells, and mast cells (Fig. S7a-d). In PN, there was increased extracellular matrix signaling and angiogenesis among FB populations, while in AD, there was an increased immune response among innate and adaptive immune cells (Fig. S7e-f, h). Mast cells in AD differed with increased expression of genes IGFLR1, PRG2, HCST, PTGDS, IFI30, ALOX5, SIGLEC6, and SLCO2B1. (Fig. S7g). Compared to PSO, lesional PN skin had increased PC1/2, mFB, SMC, FB1, END, and FB4, while PSO had increased KC, NKT, CD4pro, and Tregs (Fig. S8a-d). Expression differences existed in lesional PN skin compared to PSO with increased immune responses in myeloid lineage cells and increased ECM synthesis in FBs in PN (Fig. S8e-g). PSO skin has increased responses and development of cytotoxic and helper T cells (Fig. S8e-g). The RashX algorithm was implemented to holistically characterize PN resident memory cells compared to AD and PSO, as previously demonstrated^17^. Lesional PN skin mapped in between AD and PSO specific genes, demonstrating a mixed Th2/Th17 phenotype (Fig. 7g.) Two PN samples did align closer to AD and the Th2 axis (Fig. 7g).

**Fig. 7.**
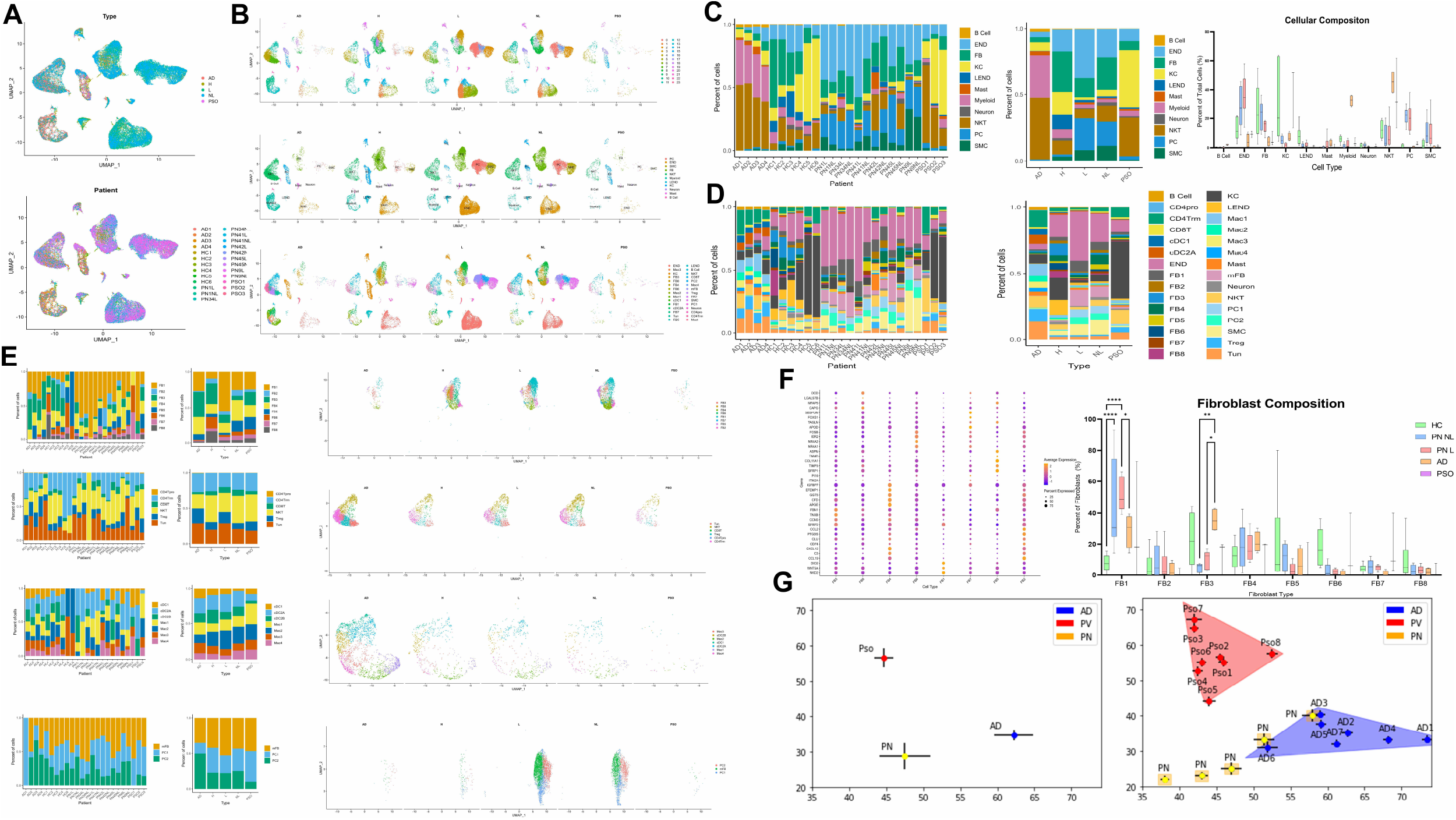
Single cell comparison of PN across AD and PSO identifies mesenchymal dominance and a mixed Th2/Th17 phenotype. **a,** UMAP of integrated cells across HC, NL, L, AD, and PSO samples colored by Type (top) and sample (bottom). **b,** UMAP of cells split per condition colored based on louvian clusters (top), annotated clusters (middle), and annotated subclusters (bottom). **c,** Stacked barplot of major clusters per sample(left), stacked barplot of average major clusters per condition (middle), and barplot of major clusters per condition. **d,** Stacked barplot of subclusters per sample (left), stacked barplot of average subclusters per condition (right). **e,** Array of stacked bar plots per sample, average stacked bar plots, and UMAPs for each major subcluster highlighting differences in subclusters within the super cluster. **f,** Dotplot displaying fibroblast subcluster defining genes using the integrated dataset, showing consistency between FB1, FB2, FB3, FB5, and FB7 with the PN cohort (left) Barplot displaying differential fibroblast composition across conditions, with FB1 increased in PN and FB3 increased in AD (right). Statistical analysis by ANOVA corrected with Tukey post hoc test with *p<0.05, ** p<0.01, ***p<0.001. **g,** Molecular classification of resident memory cells in PN L samples using RashX algorithm identifies PN as intermediate between AD and PSO Th2/Th17 expression profiles.

### PN patients have an increased risk of cancers associated with CAFs and fibroproliferative disease

Given the differences in lesional PN skin FB1 cell counts consistent across HC, AD, and PSO, we identified diseases in which this fibroblast phenotype existed. Squamous cell carcinoma is heavily associated with these CAFs, so we identified a scRNAseq dataset to map our existing clustering of FB clusters. Using this method, we found increased FB1 across lesional PN skin, with SCC having similar increased levels compared to HC, AD, and PSO (Fig. 8a). A subset of KC in lesional PN skin expressed increased midkine signaling via ligand receptor analysis compared to non-lesional skin (Fig. 8a). To characterize clinically significant risk associated with this FB population, we performed a population-level, multicenter cohort study analyzing data from TriNetX. A total of 92,965 PN patients were identified, and the majority were female (58.1%), White (65.6%), and Non-Hispanic/Latino (73.8%), with a mean age of 55.2 years (Fig 8b). PN and HC groups were balanced following 1:1 propensity score matching (Table S1). PN was associated with increased rates of all cancer outcomes, including SCCis, SCC, breast, and colorectal, with an earlier time of development (Fig. 8d-e, Table S2).

**Fig. 8.**
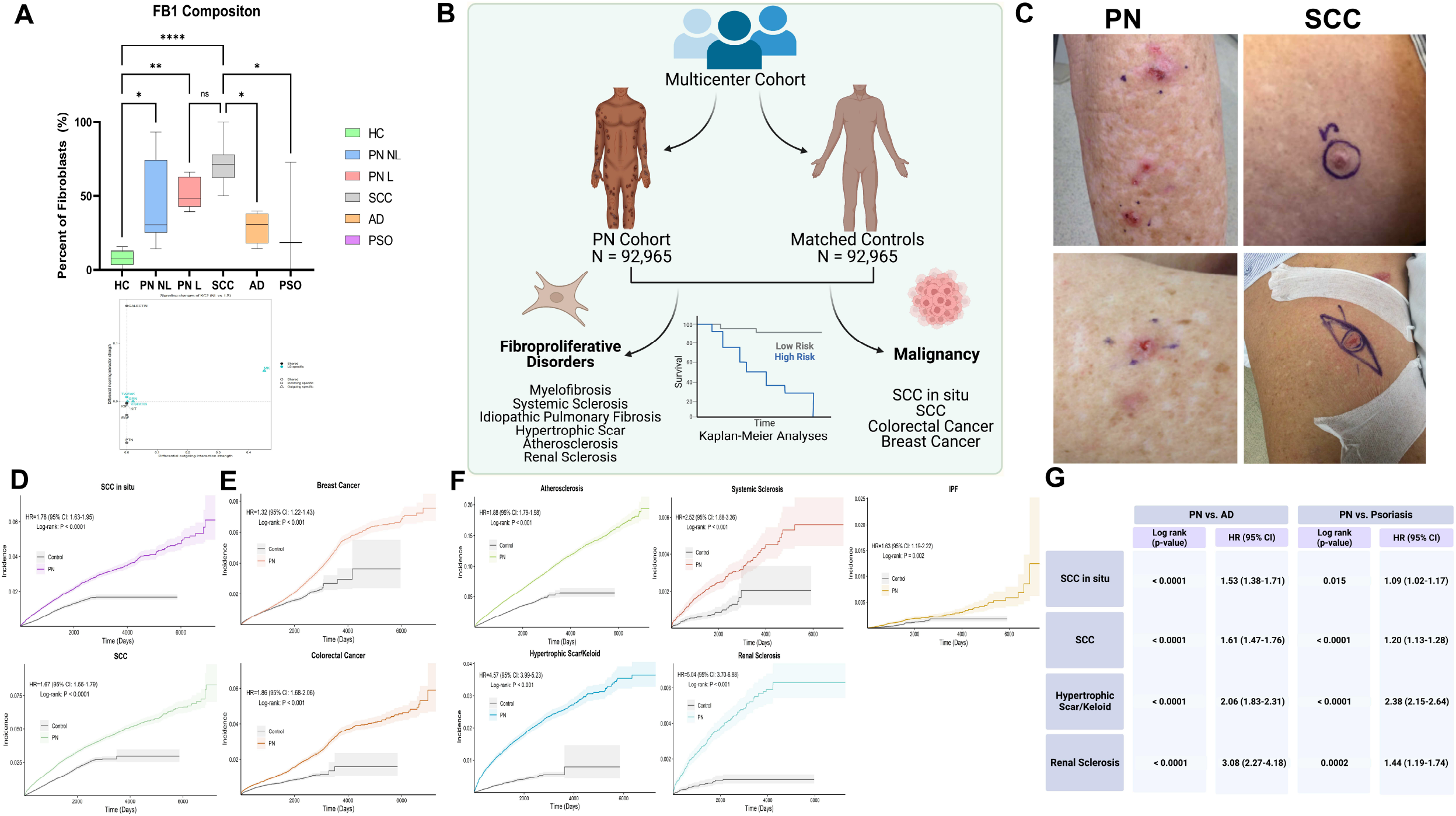
Multicenter cohort epidemiologic data analysis of diseases associated with CAF in PN reveals increased risk for SCC and fibroproliferative disease. **a,** FB1 composition across H, NL, L, SCC, AD, and PSO samples reveals increased FB1 across PN and SCC compared to HC, AD, and PSO. Statistical analysis by ANOVA corrected with Tukey post hoc test with *p<0.05, ** p<0.01, ***p<0.001 **b,** Overview graphic of multicenter cohort study comparing PN fibroblast associated malignancy and fibroproliferative disease compared to matched controls **c,** Representative images of SCC and PN nodules within the same patient **d,** Kaplan Meier curves showing increased SCC and SCCis risk for PN compared to matched HCs. **e,** Kaplan Meier curves showing increased risk for cancers reported with CAFs (breast and colorectal) in PN compared to matched HCs **f,** Kaplan Meier curves showing increased risk for fibroproliferative disease and atherosclerosis in PN compared to matched HCs **g,** Table of hazard ratios demonstrating increased risk of SCCis, SCC, hypertrophic/keloid scars, and renal sclerosis in PN compared to AD/PSO

Based on the literature, fibroproliferative disease and atherosclerosis were assessed due to their association with the FB subsets. PN was associated with increased rates of all fibroproliferative outcomes, including atherosclerosis, systemic sclerosis, idiopathic pulmonary fibrosis (IPF), hypertrophic scars, and renal sclerosis with an earlier time of development (Fig. 8f). Similarly, after adjustment for several important confounding variables in Cox proportional-hazards models, the increased risk was observed in PN patients compared to HCs for all fibroproliferative primary endpoints (Fig. 8f). Comparing PN to matched AD and PSO patients revealed increased rates of SCCis, SCC, and renal sclerosis (Fig. 8g).

## DISCUSSION

Progress in understanding the pathogenesis of PN has lagged behind other inflammatory skin diseases. Lack of studies with human patients and the absence of specific disease-tailored therapeutic targets have hampered progress. In this study, we use scRNA-seq to provide a molecular map of the multicellular landscape of PN skin. A key finding of our study is the discovery of the WNT5A+ POSTN+FB and mFB clusters that are shifted towards a CAF-like phenotype in PN skin. The real-world impact of these studies is supported by multi-center data placing PN patients at increased risk for CAF-associated cancers, highlighting the need to screen PN patients for these malignancies. These results strengthen previous single institution findings of increased malignancy risk in PN patients, in particular SCC^18^. Ligand receptor analysis of these fibroblasts also identified CAF-associated WNT5A and periostin in lesional FBs interacting with neuronal MCAM and ITGAV, suggesting a fibroblast-neuronal axis in PN. To our knowledge, this study is the first to build a single-cell atlas of PN skin, facilitating the understanding of PN pathogenesis and providing potential targets for treatment.

Overall, the cellular composition of lesional PN skin shifted towards FB and mFB populations, with increased mesenchymal proliferation and a higher mesenchymal-to-immune cell ratio compared to non-lesional PN skin. In lesional PN skin, each FB population is shifted towards this CAF-like phenotype, particularly FB1. CAFs regulate cancer metastasis through initiating the extracellular matrix and secreting growth factors, stimulating angiogenesis, modulating the immune system, and contributing to drug resistance^19,20^. FB1 cells exhibited increased expression of CAF genes (CTHRC1, NREP, WNT5A, CPXM1, GGT5, POSTN), while mFB expressed increased type I IFN and CPXM1. FB1 closely aligns with desmoplastic CAFs while mFB align with myofibroblast CAFs as previously reported across tumors^21^. This discovery of lesional fibroblasts aligning with CAFs suggests local programming in PN, which may ultimately lead to pathologic itch and proliferation^21^. WNT5A+ FB1 may promote carcinogenesis via soluble WNT5A secretion and interaction with keratinocytes, as increased proliferation in colorectal cancer from CAF WNT5A has been previously reported^22^. CPXM1 has been described as a potential pro-cancer driver in melanoma, and its expression was significantly decreased in melanoma patients responding to anti-PD1 therapy^23^. FB periostin may further augment oncogenesis in PN by promoting proliferation and invasion of cutaneous squamous cell, nasopharyngeal, and breast carcinomas^24–27^. The shift of FBs and mFBs toward a CAF-like phenotype may pose an increased risk of CAF-associated cancers for PN patients. Our multi-center analyses confirmed that PN was associated with increased rates of CAF-associated cancers, including SCC, SCCis, breast, and colorectal cancers compared to HCs, AD, and PSO. Regular screening of underlying malignancy in PN patients may improve long term outcomes. Future validation of the pro-carcinogenic potential of PN FBs is necessary via co-culturing of malignant cells and other techniques to further support the CAF phenotype.

The WNT5A+FB1 subpopulation was increased in lesional and non-lesional PN compared to HC skin, suggesting systemic disease in PN patients and priming of non-lesional skin. WNT5A is a non-canonical WNT ligand that mediates interactions among keratinocytes, immune cells, and inflammatory factors and has been implicated in inflammatory responses in PSO and skin cancer^28–30^. WNT5A in FBs regulates proliferation and provides resistance to apoptosis, a process previously shown to be pathogenic in pulmonary fibrosis^31^. Our ligand receptor analysis showed that in lesional PN skin, WNT5A was exclusively expressed by FB1 cells reacting with mesenchymal/neuronal melanoma cell adhesion molecule (MCAM). MCAM, also known as CD146, is a membrane CAM of the immunoglobulin superfamily and functions in mammalian nervous system development^32^. MCAM also facilitates nerve regeneration and regulates cell migration and proliferation, suggesting a role for a fibroblast-neuronal axis contributing to PNS hypertrophy and activation in PN^33,34^.

Increased angiogenesis markers in the endothelium of lesional skin compared to non-lesional skin suggest vascular fragility and permeability, leading to increased leukocyte extravasation and inflammatory responses in PN. Permeability may be enhanced by increased expression of LGALS in lesional endothelium, as it regulates leukocyte migration with involvement in autoimmunity, wound healing, and carcinogenesis^35,36^. Endothelial-derived SPARC and HSPG2 in lesional skin may further influence fibroblast-mediated pathology as they stimulate fibrosis and keratinocyte proliferation in systemic sclerosis and cirrhosis^37–41^. A pathogenic loop may exist with FBs stimulating angiogenesis via secreted frizzled-related protein 2 (SFRP2)^42^. Anti-SFRP2 treatment may benefit two-fold by targeting both angiogenesis-related pruritus while also inhibiting FB WNT5A^42,43^.

PN skin demonstrated pericyte to mFB transition as well as endothelial to pericyte transition. Pericyte to mFB transition can occur during microvascular damage, supporting a role for preventative vasculature directed therapy^44^. Endothelial to mesenchymal transition is an important source of CAFs and contributes to formation of aberrant tumor vessels^45,46^.

The immune cell landscape of lesional PN skin had increases in CD4+ T cells and cDC2A cells compared to non-lesional PN skin. There was also a shift towards a type I interferon response in lesional PN myeloid and mesenchymal cells. Type I interferons are involved in autoimmunity, and in PN may promote the recruitment of inflammation that drives pruritus^47^. FB-derived interferon may play a role in stimulating the immune system and promoting inflammation, all while increasing carcinogenesis risk^48^. The overall PN resident memory cell signature was identified as an intermediate between AD and PSO using the RashX algorithm^17^. This suggests a mixed Th2/Th17 response in PN, delineating it from the two diseases. Both Th2 and Th17 responses have previously been implicated in PN skin^49^. Compared to AD skin, PN lesional skin had increased mFB, PC1, FB1, and neurons, while AD had increased immune cells, including macrophages, T cells, dendritic cells, and mast cells. Compared to PSO skin, PN skin showed increased PC1/2, mFB, SMC, FB1, END, and FB4, while PSO skin showed increased KC, NKT, CD4pro, and Tregs. A mixed approach may be needed to target PN patients by fine-tuning the Th2/Th17 balance.

Finally, given the systemic dysregulation of FBs, we demonstrated that PN patients have an increased risk of fibroproliferative disease aside from CAF-associated cancers. PN FBs are prone to activation, and this may exist across organ systems, particularly in the kidney, as renal sclerosis risk was increased compared to AD/PSO. Systemic anti-fibrotic treatments for conditions such as pulmonary fibrosis may play a role in PN by modulating FB responses and preventing these long-term outcomes.

This study is limited by studying primarily the transcriptome, which does not necessarily represent protein abundance, however key findings were confirmed using protein techniques and tissue immunofluorescence staining. Future studies should include large sample sizes from diverse patient populations to allow for a comparison of racial and ethnic differences in the molecular basis of PN on the single-cell level.

Here we demonstrate that FBs from PN skin are shifted toward a CAF-like phenotype, potentially increasing the risk for developing CAF-related cancers. Clinicians should have increased suspicion for CAF-associated malignancies in PN patients and maintain a low screening threshold. The novel WNT5A+ and POSTN+ FB populations may serve as future therapeutic targets in PN.

## MATERIALS AND METHODS

### Patient Cohort and sample acquisition

This study was approved by the Johns Hopkins Institutional Review Board (IRB00119007). Adult patients with prurigo nodularis were recruited prospectively based on inclusion criteria modeled after previous clinical trials of PN, including PN diagnosed by a board-certified dermatologist, with moderate-to-severe disease as having at least 20 nodules present bilaterally on the body, and severe disease designated by WI-NRS ≥7^50^. Patients with other concomitant inflammatory skin conditions, such as psoriasis or atopic dermatitis, were excluded.

### scRNA-seq library preparation and sequencing

6 lesional and 6 non-lesional punch biopsies were processed at a single cell suspension using the 10x genomics platform.

### scRNA-seq Data Analysis

#### Data preprocessing and quality assurance

Raw sample count matrices were analyzed in R using the Seurat package^51^. Cells were filtered based on unique feature counts over 2,500 or less than 200. Cells with mitochondrial counts >5% were also filtered. Data was normalized on a log scale.

#### Cellular clustering using machine learning

Dimensional reduction using Principal Components Analysis (PCA) was performed and a principal component number of 17 was selected based on the elbow plot method. Cells were clustered using the louvain algorithm using an initial resolution of 1. Cell types were annotated based on top 10 differentially expressed genes. Subclustering was performed for major clusters with different resolutions, with the same methodology as major clusters to identify discrete subclusters. Resolution parameters were adjusted to determine the ideal resolution to prevent over or under clustering.

#### Pseudobulk RNAseq

Traditional RNAseq comparison was performed using the scRNAseq dataset using the pseudobulk method. This was performed in R using the package DESeq2^52^ to identify the top differentially regulated genes between non-lesional and lesional PN skin. Hierarchical clustering was performed to identify coregulated genes and subset samples based on composition of cell clusters such as FB1 and mFB (Figure 3h).

#### Differential cellular and transcriptomic analysis

To compare two different sample sets we utilized the case-control analysis of scRNAseq studies. This was performed in R using the package cacoa^53^. Statistical analysis to determine differential cell compositions is performed using differences in loading coefficients. Cluster free composition changes are determined by subtraction and wilcox testing. Expression differences are determined by normalized expression distance between two groups for all cell clusters. Gene ontology analysis is performed to determine top upregulated or downregulated pathways in each cellular cluster. Volcano plots determine differentially expressed genes based on predetermined logfold changes and p value thresholds.

#### Correlation analysis of cell compositions

Correlation matrices for all cell clusters were created using the Scanpy package in Python^54^.

#### Ligand Receptor Analysis

Intercellular communication pathways were analyzed using CellChat^55^ to identify differentially expressed ligands and receptors. Differential ligand receptor analysis was performed to compare non-lesional and lesional PN skin. Downstream analysis of differentially regulated pathways was performed separately on non-lesional and lesional skin to include the noncanonical WNT, periostin, and complement pathways.

#### RNA velocity and pseudotime analysis

Spliced and unspliced counts were constructed from scRNA-seq data using the velocyto^56^ pipeline (the Python script velocyto.py was run on the CellRanger output folder). RNA velocity experiments were conducted with scVelo, an integrated Scanpy package in Python, using the dynamical model to infer velocities. To perform pseudotime analyses, we used R package Monocle2^57^. After converting a Seurat integrated object into a Monocle cds object, we inferred pseudotime trajectories for FB, mFB, and PC cells.

#### Comparison to public datasets of HC, AD, PSO, SCC

Public datasets of scRNAseq data were obtained from fresh samples existing in the gene expression omnibus repository^58^. These included 6 HC samples^59^, 7 SCC samples^60^, 4 AD samples^61,62^, and 3 PSO samples^59^. HC, AD, and PSO samples were combined into a single object using Seurat and processed as above. Samples were integrated using cross dataset pairs of cells (anchors) to correct technical differences due to batch effects across studies. Subsequent analysis to identify cellular subsets and differential transcriptomic analysis was performed as above. FB subsets identified using the integrated dataset were projected onto the SCC dataset to identify similarities in CAFs. The RashX algorithm was used to compare PN to AD and PSO based on published methodology comparing disease-defining transcriptomes in resident T cells^17^.

#### Statistical Analysis

R and Graphpad Prism 9 were used for statistical analysis. When using the cacao package to compare two groups, statistical analysis was performed according to the package methods. Comparison of cell composition outside of the cacoa package was performed using nonparametric Mann Whitney test using p<0.05. DEGs between two groups for any cluster were compared using the wilcoxon rank sum test corrected using the bonferroni method on a cellular basis per sample. Significant DEG were further verified at the sample level using mean gene expression per sample and subsequent Wilcox testing performed with FDR=5%. FB subset composition comparison between HC, SCC, AD, PSO was performed using ANOVA corrected for multiple hypothesis testing with Tukey’s post hoc test. Correlation analysis was performed using the spearman r test with p<0.05 as significant.

#### Visualization

R packages as described above were used for visualization. Graph Prism 9 was used to create bar plots and correlation plots.

#### Immunofluorescent Staining

Skin samples of 5-μm thickness, formalin-fixed and paraffin embedded were deparaffinized with heat-idcued antigen retrieval using Triology buffer (Trilogy® 920P x1, Sigma-Aldrich) and blocked with DAKO protein blocking reagent (X0909, DAKO, Capinteria, CA, USA) for one hour. Primary antibodies included periostin (ab14041, Abcam; dilution, 1:400), SFRP2 (MABC5390, Millipore Sigma; dilution, 1:500), wnt5a (AF645, R&D Systems; dilution, 1:20), NKD2 (NBP2-13658, Novus Biologicals, CO, USA; dilution, 1:500), α-smooth muscle actin (ab5694, Abcam; dilution, 1:100), Apod (PA5-27386, ThermoFisher Scientific, Waltham, MA, USA; dilution, 1:500), CD52 (ab245681, Abcam; dilution, 1:200), and Tryptase (ab2378, Abcam; dilution, 1:1000). All primary antibodies were co-localized with vimentin (AF2105, R&D Systems; dilution, 1:20), incubated at 4°C overnight. Secondary antibodies were conjugated with an Alexa Fluor 488 and 568 at a dilution of 1:400, and mount with DAPI dye (Vector Laboratories, Burlingame, CA, USA). Samples were mounted with ProLong glass antifade Mountant (Invitrogen, Cat#: 36980). Images were acquired on a Leica SP8 confocal microscope (Leica Microsystems, Deerfield, IL, USA) in the Johns Hopkins Ross Fluorescence Imaging Center. The background normalized fluorescence intensity of the antibodies in the sub-epidermal dermis (fibroblast) and mast cells (CD52) was measured in arbitrary units (AU) using Image J software (NIH, Bethesda, MD, USA). One-way ANOVA was used to compare differences between PN L, PN NL and HC groups at statistical significance level of p□<□0.05.

### Epidemiologic analysis of cancer and fibroproliferative disease

We performed a population-level, multicenter cohort study analyzing data from TriNetX, a global federated health research network providing access to electronic medical records from approximately 89 million patients from 59 large healthcare organizations. We analyzed data from 2002-2021. Patients with PN were identified using *International Classification of Diseases, Tenth Revision, Clinical Modification* (ICD-10-CM) code L28.1, which has recently been validated^63^. Diagnostic codes before the introduction of ICD-10 were mapped using SNOMED-CT concepts^64^. Control patients included those without a diagnosis of PN. PN patients were matched to controls on age, sex, race, and ethnicity using 1:1 propensity score matching.

The primary outcomes were determined *a priori* based on literature and clinical judgment. These included squamous cell carcinoma (SCC), squamous cell carcinoma in situ (SCCis), systemic sclerosis (SS), myelofibrosis, idiopathic pulmonary fibrosis (IPF), hypertrophic scar/keloid, atherosclerosis, colorectal cancer (CRC), breast cancer (BCa), and renal sclerosis, which were identified using the respective ICD-10-CM codes. For each analysis, we excluded patients diagnosed with the outcome prior to the index date. We also excluded patients with a history of xeroderma pigmentosum for the SCC and SCCis analyses. We accounted for potential confounding variables by further matching of PN patients and controls. This included a history of actinic keratoses, transplantation status (heart, lung, kidney, liver, stem cell, bone marrow, and pancreas), and use of immunosuppressive medications (tacrolimus, sirolimus, cyclosporine, azathioprine, and mycophenolate) for the SCC and SCCis analyses, as well as prior history of tuberculosis, polycythemia vera, essential thrombocythemia, and antineoplastic use for myelofibrosis. We also matched on gastroesophageal reflux disease, obstructive sleep apnea, and tobacco use for the IPF analysis; tobacco use for hypertrophic scar/keloid; hypertension, hyperlipidemia, obesity, type 2 diabetes mellitus, and tobacco use for atherosclerosis; inflammatory bowel disease, tobacco use, obesity, history of colonic polyps and use of NSAIDs (Nonsteroidal anti-inflammatory drugs) for CRC; obesity, alcohol use disorder, hormone replacement therapy, and oral contraceptive use for BCa; and hypertension, smoking, and chronic kidney disease for renal sclerosis. Matching for the SS analysis was limited to the aforementioned demographic characteristics.

We performed time-to-event analyses and constructed Kaplan-Meier curves to compare diagnoses of outcomes between PN patients and controls. We conducted log-rank tests to test for differences between the two groups and performed Cox proportional-hazards regression to estimate the effect of PN on each of the outcomes. All p-values were two-sided and statistical significance was evaluated at the .05 α-level. The Benjamini-Hochberg method was applied to correct for multiple hypothesis tests.

## Supplementary Materials

Fig S1. Cellular composition and gene expression differences in lesional vs non-lesional PN

Fig S2. Myofibroblast correlations and TGFB expression differences in lesional vs non-lesional PN

Fig S3. Signaling network of PN shows increased complement network in non-lesional PN

Fig S4. Dynamic fibroblast gene expression in PN lesional skin

Fig. S5. Differential single cell landscape of lesional PN vs healthy skin.

Fig. S6. Differential single cell landscape of non-lesional PN vs healthy skin reveals systemic fibroblast dysfunction.

Fig. S7. Differential single cell landscape of lesional PN vs lesional AD reveals neural-fibroblast predominance in PN and adaptive immunity in AD.

Fig. S8. Differential single cell landscape of lesional PN vs lesional AD reveals fibroblast predominance in PN and epithelial-immune dysregulation in PSO

## Acknowledgments

Dr. Tyler Creamer and Linda Orzolek at the Johns Hopkins Single Cell & Transcriptomics Core

## Figures

**Fig S1. Cellular composition and gene expression differences in lesional vs non-lesional PN**

**a,** Barplot displaying composition of cell types in L and NL skin. **b,** Gene ontology heatmaps of top 50 pathways upregulated in PN L vs NL skin. **c,** Gene ontology heatmap of top 50 pathways downregulated in PN L vs NL skin. **d,** Volcano plot per cell cluster displaying differentially expressed genes in L vs NL skin.

**Fig S2. Myofibroblast correlations and TGFB expression differences in lesional vs non-lesional PN**

**a,** Correlation plots of L and NL FB and PC clusters against mFB composition revealing positive correlations with FB1/FB7 and a negative correlation with FB4. **b,** Split violin plots of TGFBI expression in L vs NL skin for populations FB1, PC1, PC2, SMC, and mFB revealing increased expression in L skin. All comparisons between NL and L gene expression at a cellular level represent p<0.05 using wilcoxon rank sum test.

**Fig S3. Signaling network of PN shows increased complement network in non-lesional PN**

**a,** Bar graphs displaying number of interactions (left) and the interaction strengths (right) of L and NL PN. **b,** Heatmap demonstrating the number of interactions (left) and the interaction strengths (right) of cellular clusters in PN. **c,** Heatmaps of outgoing and incoming signaling patterns in NL (top) and L (bottom) PN. **d,** Incoming interaction strength vs outgoing interaction strength in NL (top) and L (bottom) PN. **e,** Incoming (left) and outgoing (right) cluster patterns of pathways with NL represented as dots and L as squares. **f,** Outgoing (left) and Incoming (right) pathway analysis of ligand receptors in L PN skin. Cell patterns and communication patterns defined using k means clustering. Flow charts display cell group and pathway contribution to cellular and pathway patterns. Dot plots demonstrate the contribution of a particular cell type for a specified pathway. **g,** Differential incoming and outgoing strength of L vs NL PN skin for cell types FB1, FB3, and mFB revealing increased PERIOSTIN signaling throughout. **h,** NL Complement signaling overview with circle plot and heatmap demonstrating contribution of FB populations (top). Barplots and violin plots reveal interactions of ligands C3 from FB populations and receptors ITGAX, TGAM, and C3AR1 on cDC and immune populations (middle). Heatmap of sender, receiver, mediator, and influencer categories demonstrating for NL complement signaling (bottom).

**Fig S4. Dynamic fibroblast gene expression in PN lesional skin**

a, Visualization of dynamics of select genes: ratio of unspliced to spliced transcripts (left), RNA velocity (middle), and expression values (right). b, Heatmap of different blocks of DEGs along the pseudotime trajectory for FB lesional, FB non-lesional, FB1 lesional, and FB1 non-lesional skin.

**Fig. S5. Differential single cell landscape of lesional PN vs healthy skin**. **a,** Compositional analysis of cell clusters showing differential cell loading coefficients of FB and PCs in L compared to HC skin. Statistical analysis done via Wilcoxon rank sum test with p values shown in adjacent bar graph. **b,** Barplot displaying composition of cell types in L and HC skin. **c,** Hierarchical representation of compositional changes in L and HC skin. **d,** cluster free compositional changes based on subtraction (top) and wilcox testing (bottom). **e,** Expression differences calculated using normalized expression distance between L and HC skin for all cell clusters. **f,** cluster free expression shifts calculated by proportion change (left) and separability z score (right). **g,** Gene ontology heatmaps of top 50 pathways upregulated or downregulated in PN L vs HC skin. **h,** Volcano plot per cell cluster displaying differentially expressed genes in L vs HC skin.

**Fig. S6. Differential single cell landscape of non-lesional PN vs healthy skin reveals systemic fibroblast dysfunction**. **a,** Compositional analysis of cell clusters showing differential cell loading coefficients of FB and PCs in NL compared to HC skin. **b,** Barplot displaying composition of cell types in NL and HC skin. Statistical analysis done via Wilcoxon rank sum test with p values shown in adjacent bar graph. **c,** Hierarchical representation of compositional changes in NL and HC skin. **d,** cluster free compositional changes based on subtraction (top) and wilcox testing (bottom). **e,** Expression differences calculated using normalized expression distance between NL and HC skin for all cell clusters. **f,** Gene ontology heatmaps of top 50 pathways upregulated or downregulated in PN L vs HC skin. **g,** Dotplot displaying DEGs for FB1, FB3, and FB7 NL populations displaying a systemic dysregulation compared to HC skin. **h,** Dotplot of DEGs for NL skin neurons compared to HC skin neurons. **i,** Split violin plot displaying increased pruritogenic IL31 expression in CD4Trm in NL skin compared to HC. **j,** Volcano plot per cell cluster displaying differentially expressed genes in NL vs HC skin. All dot plot comparisons between H and NL gene expression at a cellular level represent p<0.05 using wilcoxon rank sum test. All asterisks represent multiple t tests at a sample level using mean cellular expression, with *p<0.5 and ** p<0.01.

**Fig. S7. Differential single cell landscape of lesional PN vs lesional AD reveals neural-fibroblast predominance in PN and adaptive immunity in AD. a,** Compositional analysis of cell clusters showing differential cell loading coefficients of FB in L compared to immune cells in AD skin. **b,** Barplot displaying composition of cell types in L and AD skin. Statistical analysis done via Wilcoxon rank sum test with p values shown in adjacent bar graph. **c,** Hierarchical representation of compositional changes in L and AD skin. **d,** cluster free compositional changes based on subtraction (top) and wilcox testing (bottom). **e,** Expression differences calculated using normalized expression distance between L and AD skin for all cell clusters. **f,** Gene ontology heatmaps of top 50 pathways upregulated or downregulated in PN L vs AD skin. **g,** Dotplot displaying DEGs for Mast cells in L compared to AD. **h,** Volcano plot per cell cluster displaying differentially expressed genes in L vs AD skin. All dot plot comparisons between H and NL gene expression at a cellular level represent p<0.05 using wilcoxon rank sum test. All asterisks represent multiple t tests at a sample level using mean cellular expression, with *p<0.5 and ** p<0.01.

**Fig. S8. Differential single cell landscape of lesional PN vs lesional AD reveals fibroblast predominance in PN and epithelial-immune dysregulation in PSO. a,** Compositional analysis of cell clusters showing differential cell loading coefficients of FB in L compared to KC and T cells in PSO skin. **b,** Barplot displaying composition of cell types in L and PSO skin. Statistical analysis done via Wilcoxon rank sum test with p values shown in adjacent bar graph. **c,** Hierarchical representation of compositional changes in L and PSO skin. **d,** cluster free compositional changes based on subtraction (top) and wilcox testing (bottom). **e,** Expression differences calculated using normalized expression distance between L and PSO skin for all cell clusters. **f,** Gene ontology heatmaps of top 50 pathways upregulated or downregulated in PN L vs PSO skin. **g,** Split violin plots of DEGs in CD4Trm (IL13, IL17A) revealing Th2 shift in PN compared to PSO. Statistical analysis of PSO vs PN L gene expression at a cellular level represent p<0.05 using wilcoxon rank sum test. **h,** Volcano plot per cell cluster displaying differentially expressed genes in L vs PSO skin.

