## Supplementary Materials for "Single-cell RNA sequencing reveals dysregulated fibroblast subclusters in prurigo nodularis"

**Table S1. Baseline demographic characteristics.** Demographics of prurigo nodularis (PN) and control patients before and after propensity score matching.

| | **Before matching** | | | **After matching** | | |
|---|---|---|---|---|---|---|
| | PN (n = 95,813) | Control (n = 542,587) | *P-val* | PN (n = 92,965) | Control (n = 92,965) | *P-val* |
| Age, mean ± SD | 55.2 ± 15.2 | 50.9 ± 17.6 | < 0.0001 | 55.2 ± 15.2 | 55.2 ± 15.4 | 0.004 |
| Sex | | | | | | |
| Female, % | 57.8% | 58.2% | 0.03 | 58.1% | 58.3% | 0.30 |
| Male, % | 42.2% | 41.8% | 0.02 | 41.9% | 41.6% | 0.26 |
| Race | | | | | | |
| White, % | 65.1% | 76.0% | < 0.001 | 65.6% | 65.6% | 0.81 |
| Black, % | 16.8% | 11.6% | < 0.001 | 16.9% | 16.6% | 0.16 |
| Asian, % | 2.7% | 2.5% | < 0.001 | 2.7% | 2.8% | 0.28 |
| American Indian or Alaska Native, % | 0.4% | 0.3% | 0.001 | 0.4% | 0.4% | 0.40 |
| Native Hawaiian or Other Pacific Islander, % | 0.1% | 0.1% | 0.02 | 0.1% | 0.1% | 0.76 |
| Unknown, % | 14.9% | 9.5% | < 0.001 | 14.3% | 14.5% | 0.14 |
| Ethnicity | | | | | | |
| Hispanic or Latino, % | 6.3% | 10.8% | < 0.001 | 6.4% | 6.5% | 0.58 |
| Not Hispanic or Latino, % | 73.6% | 74.2% | < 0.001 | 73.8% | 74.7% | < 0.001 |
| Unknown, % | 20.1% | 15.0% | < 0.001 | 19.8% | 18.8% | < 0.001 |

**Table S2. Incidence comparison of CAF related malignancies and fibroproliferative disease in PN vs matched controls**

| | Prurigo Nodularis | | | Matched Controls | | | |
|---|---|---|---|---|---|---|---|
| | Total patients, n | Patients with outcome, n | IR (%) | Total patients, n | Patients with outcome, n | IR (%) | HR (95% CI) |
| **SCC** | 89,983 | 2,390 | 2.66 | 91,938 | 1,175 | 1.28 | 1.67 (1.55-1.79) |
| **SCCis** | 90,899 | 1,622 | 1.78 | 92,489 | 734 | 0.79 | 1.78 (1.63-1.95) |
| **Systemic Sclerosis** | 95,478 | 200 | 0.21 | 95,744 | 62 | 0.07 | 2.52 (1.88-3.36) |
| **IPF** | 95,498 | 144 | 0.15 | 95,581 | 61 | 0.06 | 1.63 (1.19-2.22) |
| **Hypertrophic Scar/Keloid** | 94,136 | 1,383 | 1.47 | 95,596 | 254 | 0.27 | 4.57 (3.99-5.23) |
| **Atherosclerosis** | 91,233 | 5,089 | 5.58 | 93,824 | 2,069 | 2.21 | 1.88 (1.79-1.98) |
| **Colorectal Cancer** | 92,835 | 1,402 | 1.51 | 93,211 | 529 | 0.57 | 1.86 (1.68-2.06) |
| **Breast Cancer** | 92,196 | 1,918 | 2.08 | 93,102 | 990 | 1.06 | 1.32 (1.22-1.43) |
| **Renal Sclerosis** | 95,324 | 287 | 0.30 | 95,587 | 47 | 0.05 | 5.04 (3.70-6.88) |

IR, Incidence Rate
HR, Hazard ratio
CI, Confidence Interval

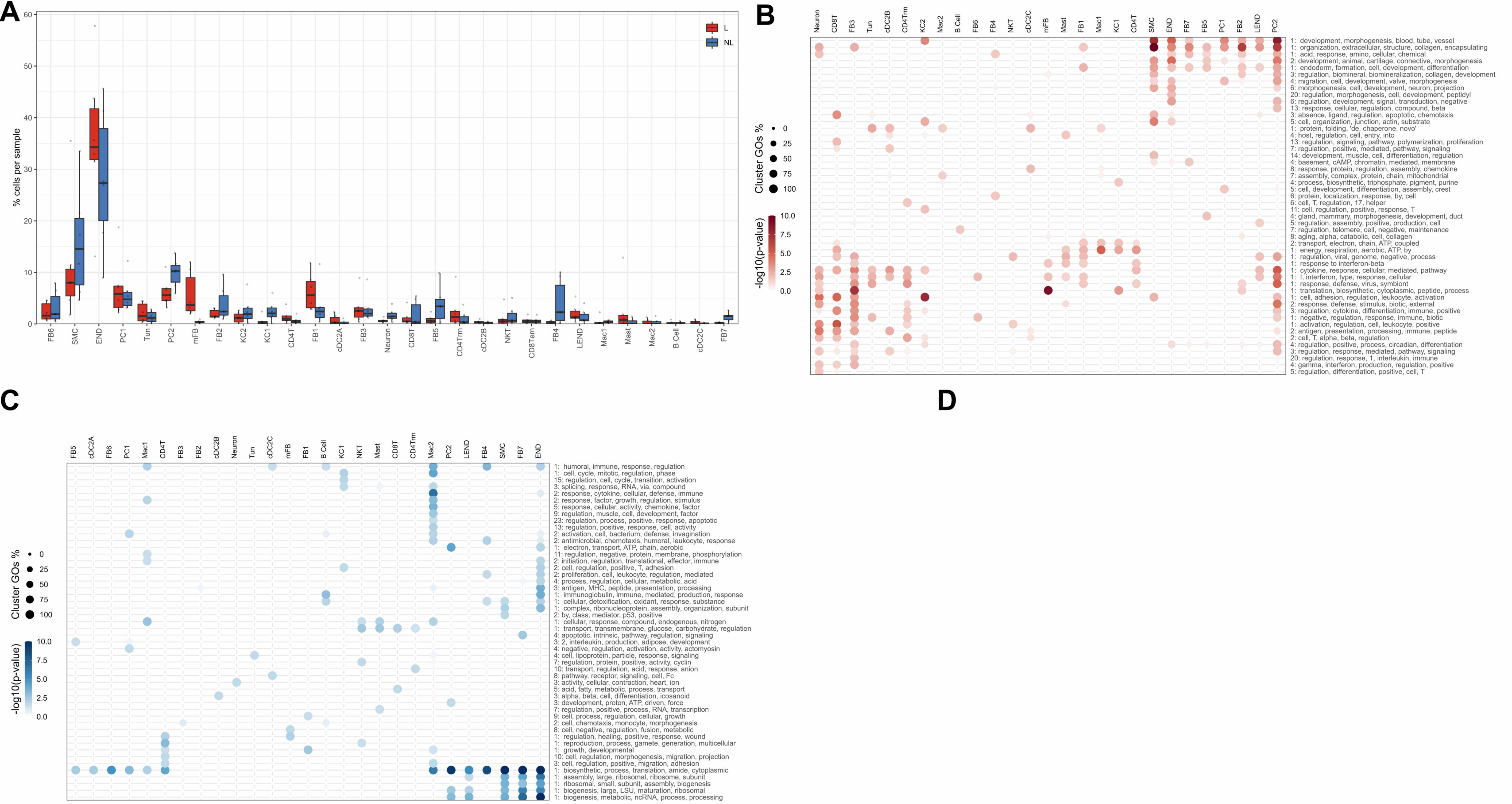
A
B
C
D
% cells per sample
L
NL
FB6
SMC
END
PC1
Tun
PC2
mFB
FB2
KC2
KC1
CD4T
FB1
cDC2A
FB3
Neuron
CD8T
FB5
CD4Trm
cDC2B
NKT
CD8Tem
FB4
LEND
Mac1
Mast
Mac2
B Cell
cDC2C
FB7
Cluster GOs %
-log10(p-value)
1: development, morphogenesis, blood, tube, vessel
1: organization, extracellular, structure, collagen, encapsulating
1: acid, response, amino, cellular, chemical
2: development, animal, cartilage, connective, morphogenesis
1: endoderm, formation, cell, development, differentiation
3: regulation, biomineral, biomineralization, collagen, development
4: migration, cell, development, valve, morphogenesis
6: morphogenesis, cell, development, neuron, projection
20: regulation, morphogenesis, cell, development, peptidyl
6: regulation, development, signal, transduction, negative
13: response, cellular, regulation, compound, beta
3: absence, ligand, regulation, apoptotic, chemotaxis
5: cell, organization, junction, actin, substrate
1: protein, folding, 'de, chaperone, novo'
4: host, regulation, cell, entry, into
13: regulation, signaling, pathway, polymerization, proliferation
7: regulation, positive, mediated, pathway, signaling
14: development, muscle, cell, differentiation, regulation
4: basement, cAMP, chromatin, mediated, membrane
8: response, protein, regulation, assembly, chemokine
7: assembly, complex, protein, chain, mitochondrial
4: process, biosynthetic, triphosphate, pigment, purine
5: cell, development, differentiation, assembly, crest
6: protein, localization, response, by, cell
6: cell, T, regulation, 17, helper
11: cell, regulation, positive, response, T
4: gland, mammary, morphogenesis, development, duct
5: regulation, assembly, positive, production, cell
7: regulation, telomere, cell, negative, maintenance
8: aging, alpha, catabolic, cell, collagen
2: transport, electron, chain, ATP, coupled
1: energy, respiration, aerobic, ATP, by
1: regulation, viral, genome, negative, process
1: response to interferon-beta
1: cytokine, response, cellular, mediated, pathway
1: I, interferon, type, response, cellular
1: response, defense, virus, symbiont
1: translation, biosynthetic, cytoplasmic, peptide, process
1: cell, adhesion, regulation, leukocyte, activation
2: response, defense, stimulus, biotic, external
3: regulation, cytokine, differentiation, immune, positive
1: negative, regulation, response, immune, biotic
1: activation, regulation, cell, leukocyte, positive
2: antigen, presentation, processing, immune, peptide
2: cell, T, alpha, beta, regulation
4: regulation, positive, process, circadian, differentiation
3: regulation, response, mediated, pathway, signaling
20: regulation, response, 1, interleukin, immune
4: gamma, interferon, production, regulation, positive
5: regulation, differentiation, positive, cell, T
1: humoral, immune, response, regulation
1: cell, cycle, mitotic, regulation, phase
15: regulation, cell, cycle, transition, activation
3: splicing, response, RNA, via, compound
2: response, cytokine, cellular, defense, immune
2: response, factor, growth, regulation, stimulus
5: response, cellular, activity, chemokine, factor
9: regulation, muscle, cell, development, factor
23: regulation, process, positive, response, apoptotic
13: regulation, positive, response, cell, activity
2: activation, cell, bacterium, defense, invagination
2: antimicrobial, chemotaxis, humoral, leukocyte, response
1: electron, transport, ATP, chain, aerobic
11: regulation, negative, protein, membrane, phosphorylation
2: initiation, regulation, translational, effector, immune
2: cell, regulation, positive, T, adhesion
2: proliferation, cell, leukocyte, regulation, mediated
4: process, regulation, cellular, metabolic, acid
3: antigen, MHC, peptide, presentation, processing
1: immunoglobulin, immune, mediated, production, response
1: cellular, detoxification, oxidant, response, substance
1: complex, ribonucleoprotein, assembly, organization, subunit
2: by, class, mediator, p53, positive
1: cellular, response, compound, endogenous, nitrogen
1: transport, transmembrane, glucose, carbohydrate, regulation
4: apoptotic, intrinsic, pathway, regulation, signaling
3: 2, interleukin, production, adipose, development
4: negative, regulation, activation, activity, actomyosin
4: cell, lipoprotein, particle, response, signaling
7: regulation, protein, positive, activity, cyclin
10: transport, regulation, acid, response, anion
8: pathway, receptor, signaling, cell, Fc
3: activity, cellular, contraction, heart, ion
5: acid, fatty, metabolic, process, transport
3: alpha, beta, cell, differentiation, icosanoid
3: development, proton, ATP, driven, force
7: regulation, positive, process, RNA, transcription
9: cell, process, regulation, cellular, growth
2: cell, chemotaxis, monocyte, regulation, morphogenesis
8: cell, negative, regulation, fusion, metabolic
1: regulation, healing, positive, response, wound
1: reproduction, process, gamete, generation, multicellular
1: growth, developmental
10: cell, regulation, morphogenesis, migration, projection
3: cell, regulation, positive, migration, adhesion
1: biosynthetic, process, translation, amide, cytoplasmic
1: assembly, large, ribosomal, ribosome, subunit
1: ribosomal, small, subunit, assembly, biogenesis
1: biogenesis, large, LSU, maturation, ribosomal
1: biogenesis, metabolic, ncRNA, process, processing

**Fig S1. Cellular composition and gene expression differences in lesional vs non-lesional PN**

**a,** Barplot displaying composition of cell types in L and NL skin. **b,** Gene ontology heatmaps of top 50 pathways upregulated in PN L vs NL skin. **c,** Gene ontology heatmap of top 50 pathways downregulated in PN L vs NL skin. **d,** Volcano plot per cell cluster displaying differentially expressed genes in L vs NL skin.

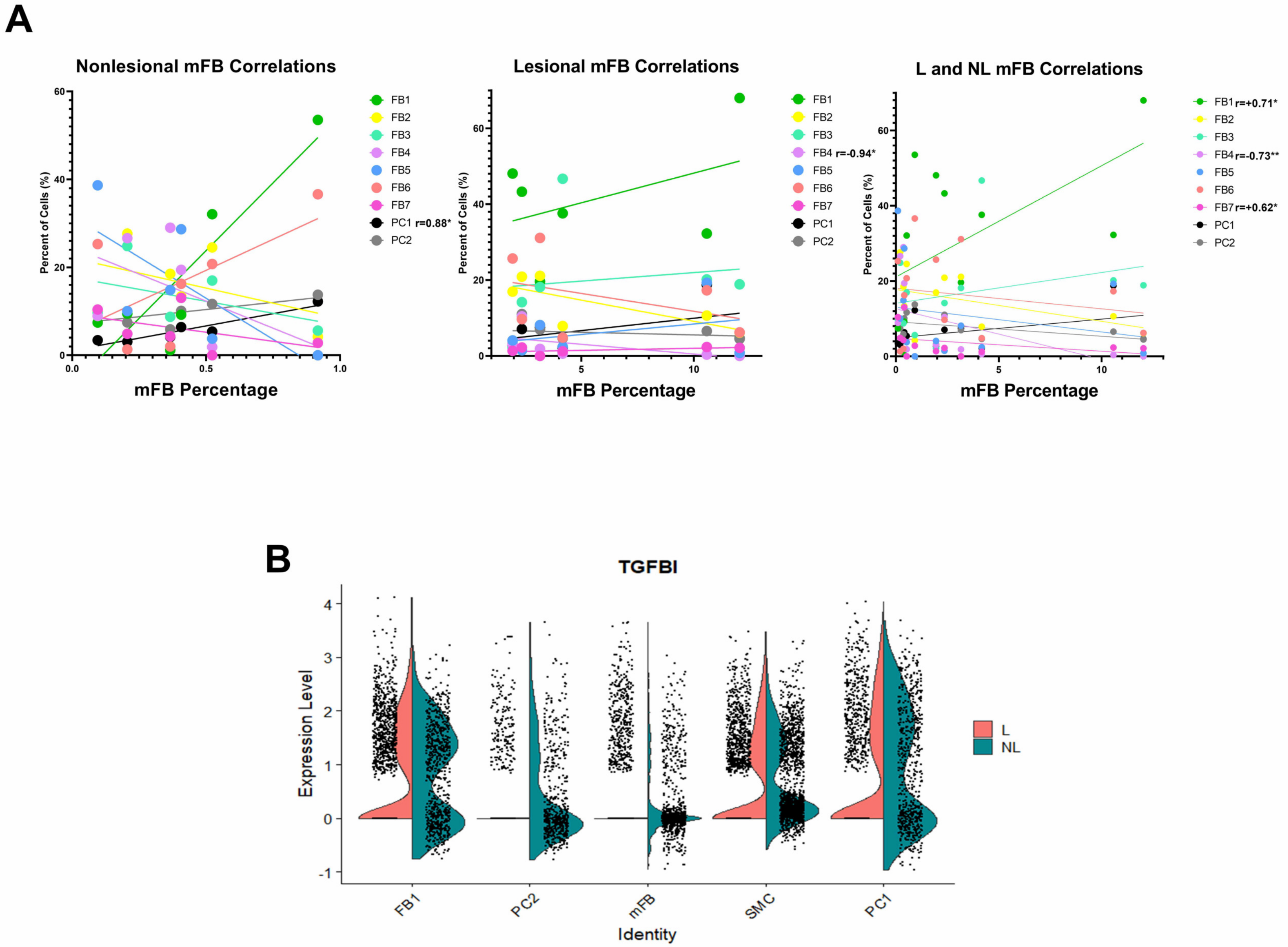
A
Nonlesional mFB Correlations
Lesional mFB Correlations
L and NL mFB Correlations
Percent of Cells (%)
mFB Percentage
FB1
FB2
FB3
FB4
FB5
FB6
FB7
PC1
PC2
r=0.88*
r=-0.94*
r=+0.71*
r=-0.73**
r=+0.62*
B
TGFBI
Expression Level
Identity
FB1
PC2
mFB
SMC
PC1
L
NL

**Fig S2. Myofibroblast correlations and TGFB expression differences in lesional vs non-lesional PN**

**a,** Correlation plots of L and NL FB and PC clusters against mFB composition revealing positive correlations with FB1/FB7 and a negative correlation with FB4. **b,** Split violin plots of TGFBI expression in L vs NL skin for populations FB1, PC1, PC2, SMC, and mFB revealing increased expression in L skin. All comparisons between NL and L gene expression at a cellular level represent $p<0.05$ using wilcoxon rank sum test.

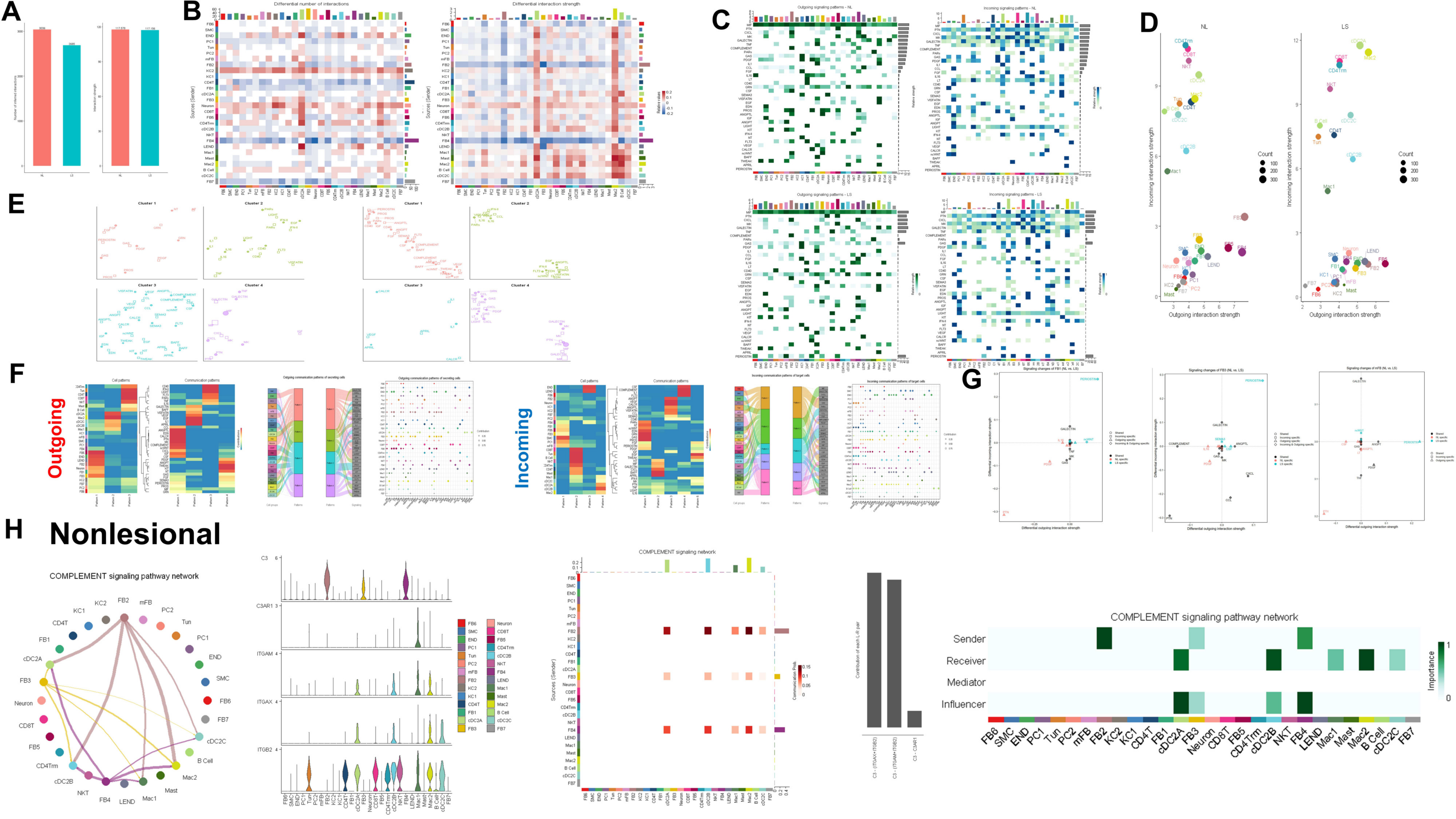
A
B
Differential number of interactions
Differential interaction strength
C
Outgoing signaling patterns - NL
Incoming signaling patterns - NL
Outgoing signaling patterns - LS
Incoming signaling patterns - LS
D
NL
LS
Incoming interaction strength
Outgoing interaction strength
Count
E
Cluster 1
Cluster 2
Cluster 3
Cluster 4
F
Outgoing
Incoming
Cell patterns
Communication patterns
G
H
Nonlesional
COMPLEMENT signaling pathway network
COMPLEMENT signaling network
Sources (Sender)
Contribution of each L-R pair
Sender
Receiver
Mediator
Influencer
Importance

**Fig S3. Signaling network of PN shows increased complement network in non-lesional PN**

**a,** Bar graphs displaying number of interactions (left) and the interaction strengths (right) of L and NL PN. **b,** Heatmap demonstrating the number of interactions (left) and the interaction strengths (right) of cellular clusters in PN. **c,** Heatmaps of outgoing and incoming signaling patterns in NL (top) and L (bottom) PN. **d,** Incoming interaction strength vs outgoing interaction strength in NL (top) and L (bottom) PN. **e,** Incoming (left) and outgoing (right) cluster patterns of pathways with NL represented as dots and L as squares. **f,** Outgoing (left) and Incoming (right) pathway analysis of ligand receptors in L PN skin. Cell patterns and communication patterns defined using k means clustering. Flow charts display cell group and pathway contribution to cellular and pathway patterns. Dot plots demonstrate the contribution of a particular cell type for a specified pathway. **g,** Differential incoming and outgoing strength of L vs NL PN skin for cell types FB1, FB3, and mFB revealing increased PERIOSTIN signaling throughout. **h,** NL Complement signaling overview with circle plot and heatmap demonstrating contribution of FB populations (top). Barplots and violin plots reveal interactions of ligands C3 from FB populations and receptors ITGAX, TGAM, and C3AR1 on cDC and immune populations (middle). Heatmap of sender, receiver, mediator, and influencer categories demonstrating for NL complement signaling (bottom).

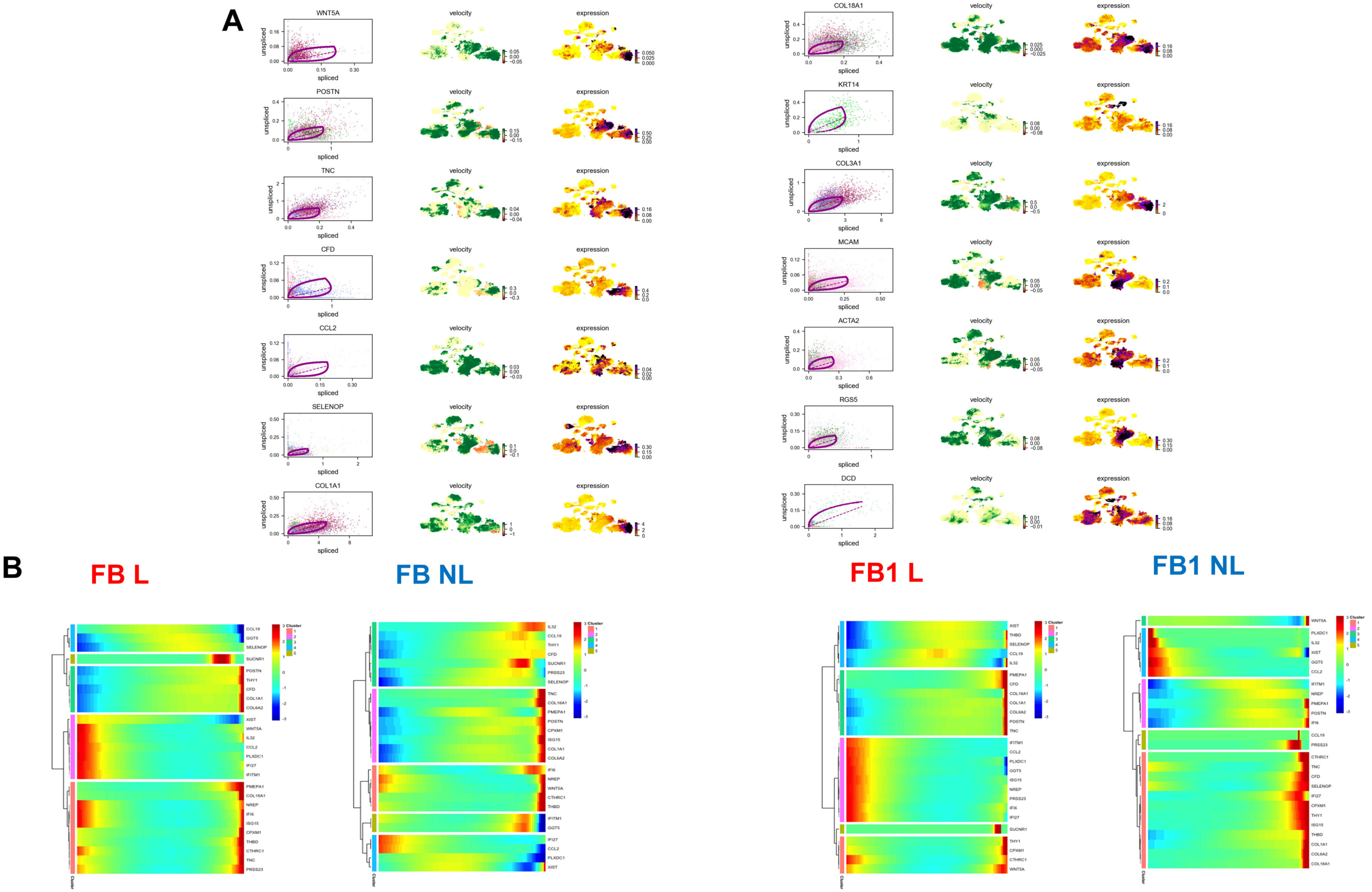

A
WNT5A
POSTN
TNC
CFD
CCL2
SELENOP
COL1A1
COL18A1
KRT14
COL3A1
MCAM
ACTA2
RGS5
DCD
velocity
expression
unspliced
spliced
B
FB L
FB NL
FB1 L
FB1 NL
Cluster

**Fig S4. Dynamic fibroblast gene expression in PN lesional skin**

a**,** Visualization of dynamics of select genes: ratio of unspliced to spliced transcripts (left), RNA velocity (middle), and expression values (right). b**,** Heatmap of different blocks of DEGs along the pseudotime trajectory for FB lesional, FB non-lesional, FB1 lesional, and FB1 non-lesional skin.

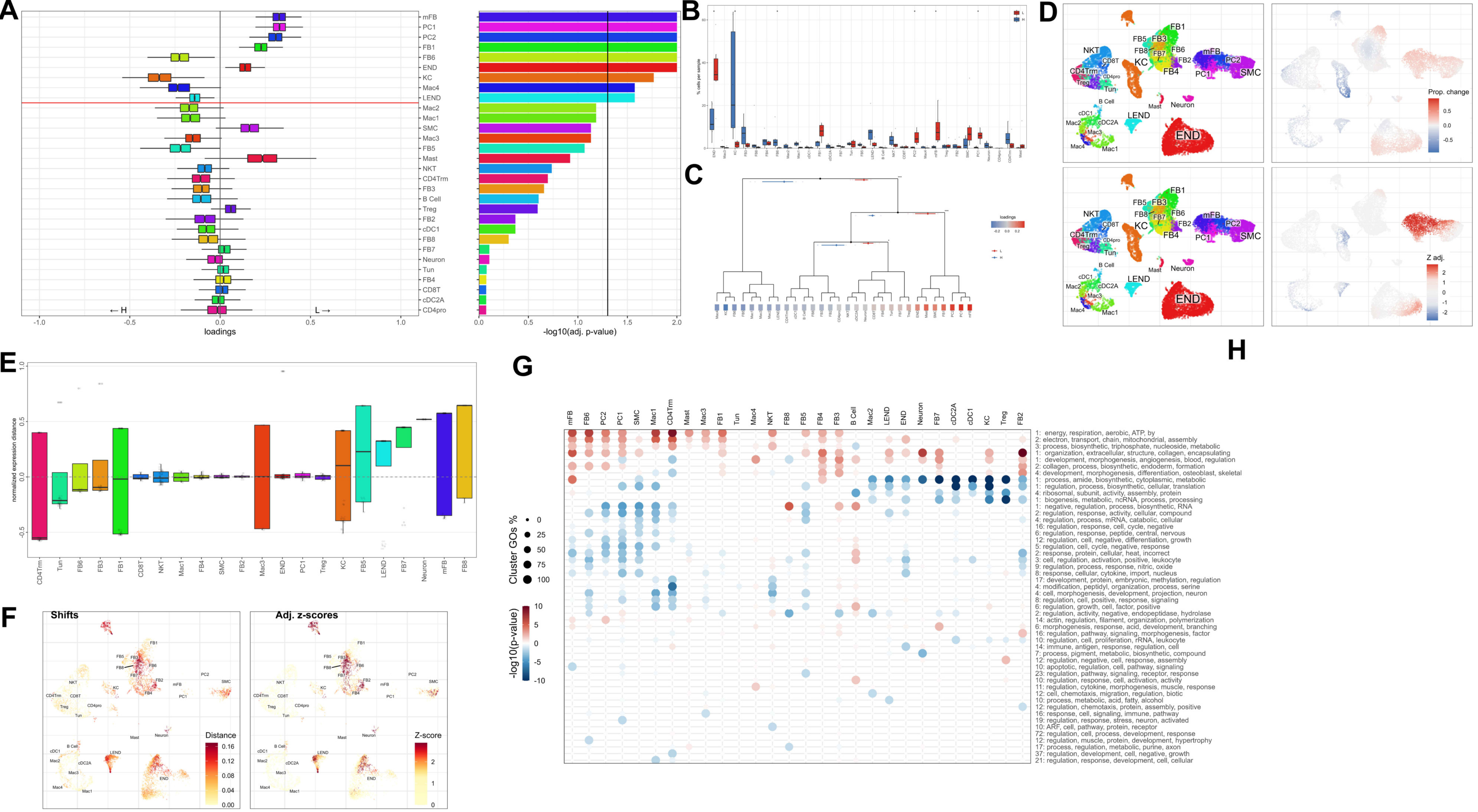

A
mFB
PC1
PC2
FB1
FB6
END
KC
Mac4
LEND
Mac2
Mac1
SMC
Mac3
FB5
Mast
NKT
CD4Trm
FB3
B Cell
Treg
FB2
cDC1
FB8
FB7
Neuron
Tun
FB4
CD8T
cDC2A
CD4pro
← H
L →
loadings
-log10(adj. p-value)
B
% cells per sample
C
loadings
D
Prop. change
Z adj.
E
normalized expression distance
F
Shifts
Adj. z-scores
Distance
Z-score
G
H
Cluster GOs %
-log10(p-value)
1: energy, respiration, aerobic, ATP, by
2: electron, transport, chain, mitochondrial, assembly
3: process, biosynthetic, triphosphate, nucleoside, metabolic
1: organization, extracellular, structure, collagen, encapsulating
1: development, morphogenesis, angiogenesis, blood, regulation
2: collagen, process, biosynthetic, endoderm, formation
4: development, morphogenesis, differentiation, osteoblast, skeletal
1: process, amide, biosynthetic, cytoplasmic, metabolic
1: regulation, process, biosynthetic, cellular, translation
4: ribosomal, subunit, activity, assembly, protein
1: biogenesis, metabolic, ncRNA, process, processing
1: negative, regulation, process, biosynthetic, RNA
2: regulation, response, activity, cellular, compound
4: regulation, process, mRNA, catabolic, cellular
16: regulation, response, cell, cycle, negative
6: regulation, response, peptide, central, nervous
12: regulation, cell, negative, differentiation, growth
5: regulation, cell, cycle, negative, response
2: response, protein, cellular, heat, incorrect
3: cell, regulation, activation, positive, leukocyte
9: regulation, process, response, nitric, oxide
8: response, cellular, cytokine, import, nucleus
17: development, protein, embryonic, methylation, regulation
4: modification, peptidyl, organization, process, serine
4: cell, morphogenesis, development, projection, neuron
8: regulation, cell, positive, response, signaling
6: regulation, growth, cell, factor, positive
2: regulation, activity, negative, endopeptidase, hydrolase
14: actin, regulation, filament, organization, polymerization
6: morphogenesis, response, acid, development, branching
16: regulation, pathway, signaling, morphogenesis, factor
10: regulation, cell, proliferation, rRNA, leukocyte
14: immune, antigen, response, regulation, cell
7: process, pigment, metabolic, biosynthetic, compound
12: regulation, negative, cell, response, assembly
10: apoptotic, regulation, cell, pathway, signaling
23: regulation, pathway, signaling, receptor, response
10: regulation, response, cell, activation, activity
11: regulation, cytokine, morphogenesis, muscle, response
12: cell, chemotaxis, migration, regulation, biotic
10: process, metabolic, acid, fatty, alcohol
12: regulation, chemotaxis, protein, assembly, positive
16: response, cell, signaling, immune, pathway
19: regulation, response, stress, neuron, activated
10: ARF, cell, pathway, protein, receptor
72: regulation, cell, process, development, response
12: regulation, muscle, protein, development, hypertrophy
17: process, regulation, metabolic, purine, axon
37: regulation, development, cell, negative, growth
21: regulation, response, development, cell, cellular

**Fig. S5. Differential single cell landscape of lesional PN vs healthy skin**. **a,** Compositional analysis of cell clusters showing differential cell loading coefficients of FB and PCs in L compared to HC skin. Statistical analysis done via Wilcoxon rank sum test with p values shown in adjacent bar graph. **b,** Barplot displaying composition of cell types in L and HC skin. **c,** Hierarchical representation of compositional changes in L and HC skin. **d,** cluster free compositional changes based on subtraction (top) and wilcox testing (bottom). **e,** Expression differences calculated using normalized expression distance between L and HC skin for all cell clusters. **f,** cluster free expression shifts calculated by proportion change (left) and separability z score (right). **g,** Gene ontology heatmaps of top 50 pathways upregulated or downregulated in PN L vs HC skin.

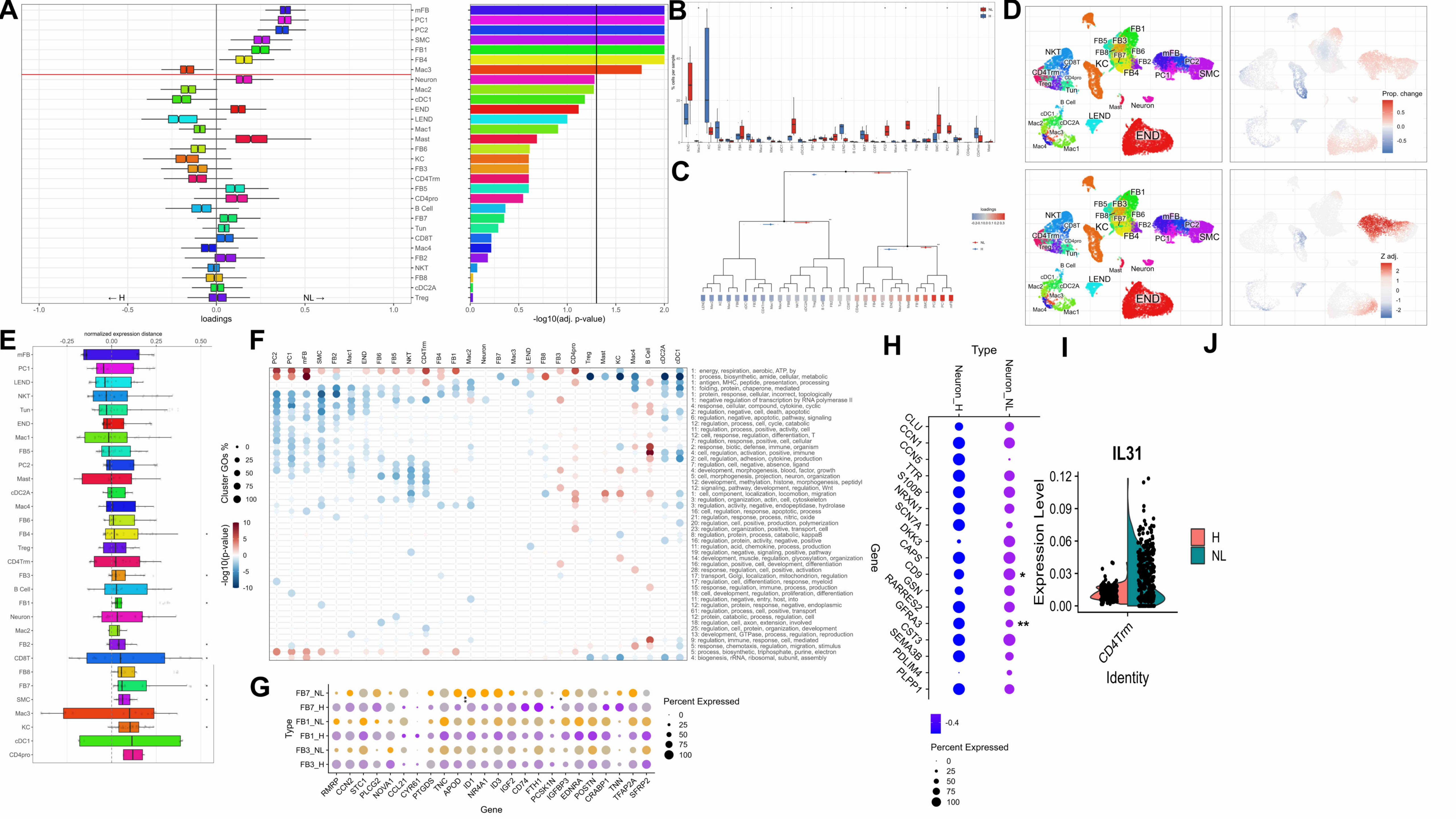
A
mFB
PC1
PC2
SMC
FB1
FB4
Mac3
Neuron
Mac2
cDC1
END
LEND
Mac1
Mast
FB6
KC
FB3
CD4Trm
FB5
CD4pro
B Cell
FB7
Tun
CD8T
Mac4
FB2
NKT
FB8
cDC2A
Treg
← H
NL →
-1.0
-0.5
0.0
0.5
1.0
loadings
0.0
0.5
1.0
1.5
2.0
-log10(adj. p-value)
B
NL
H
% cells per sample
C
loadings
NL
H
D
Prop. change
0.5
0.0
-0.5
Z adj.
2
1
0
-1
-2
FB1
FB5
FB3
FB8
FB7
FB6
FB2
FB4
mFB
PC2
PC1
SMC
NKT
CD8T
KC
CD4Trm
CD4pro
Treg
Tun
B Cell
cDC1
cDC2A
Mac2
Mac3
Mac4
Mac1
Mast
Neuron
LEND
END
E
normalized expression distance
-0.25
0.00
0.25
0.50
mFB
PC1
LEND
NKT
Tun
END
Mac1
FB5
PC2
Mast
cDC2A
Mac4
FB6
FB4
Treg
CD4Trm
FB3
B Cell
FB1
Neuron
Mac2
FB2
CD8T
FB8
FB7
SMC
Mac3
KC
cDC1
CD4pro
F
PC2
PC1
mFB
SMC
FB2
Mac1
END
FB6
FB5
NKT
CD4Trm
FB4
FB1
Mac2
Neuron
FB7
Mac3
LEND
FB8
FB3
CD4pro
Treg
Mast
KC
Mac4
B Cell
cDC2A
cDC1
Cluster GOs %
0
25
50
75
100
-log10(p-value)
10
5
0
-5
-10
1: energy, respiration, aerobic, ATP, by
1: process, biosynthetic, amide, cellular, metabolic
1: antigen, MHC, peptide, presentation, processing
1: folding, protein, chaperone, mediated
1: protein, response, cellular, incorrect, topologically
1: negative regulation of transcription by RNA polymerase II
4: response, cellular, compound, cytokine, cyclic
2: regulation, negative, cell, death, apoptotic
6: regulation, negative, apoptotic, pathway, signaling
12: regulation, process, cell, cycle, catabolic
11: regulation, process, positive, activity, cell
12: cell, response, regulation, differentiation, T
7: regulation, response, positive, cell, cellular
2: response, biotic, defense, immune, organism
4: cell, regulation, activation, positive, immune
2: cell, regulation, adhesion, cytokine, production
7: regulation, cell, negative, absence, ligand
4: development, morphogenesis, blood, factor, growth
5: cell, morphogenesis, projection, neuron, organization
12: development, methylation, histone, morphogenesis, peptidyl
12: signaling, pathway, development, regulation, Wnt
1: cell, component, localization, locomotion, migration
3: regulation, organization, actin, cell, cytoskeleton
3: regulation, activity, negative, endopeptidase, hydrolase
16: cell, regulation, response, apoptotic, process
21: regulation, response, process, nitric, oxide
20: regulation, cell, positive, production, polymerization
23: regulation, organization, positive, transport, cell
8: regulation, protein, process, catabolic, kappaB
16: regulation, protein, activity, negative, positive
11: regulation, acid, chemokine, process, production
19: regulation, negative, signaling, positive, pathway
14: development, muscle, regulation, glycosylation, organization
16: regulation, positive, cell, development, differentiation
28: response, regulation, cell, positive, activation
17: transport, Golgi, localization, mitochondrion, regulation
17: regulation, cell, differentiation, response, myeloid
15: response, regulation, immune, process, production
18: cell, development, regulation, proliferation, differentiation
11: regulation, response, entry, host, into
12: regulation, protein, response, negative, endoplasmic
61: regulation, process, cell, positive, transport
12: protein, catabolic, process, regulation, cell
18: regulation, cell, axon, extension, involved
25: regulation, cell, protein, organization, development
13: development, GTPase, process, regulation, reproduction
9: regulation, immune, response, cell, mediated
5: response, chemotaxis, regulation, migration, stimulus
5: process, biosynthetic, triphosphate, purine, electron
4: biogenesis, rRNA, ribosomal, subunit, assembly
G
FB7_NL
FB7_H
FB1_NL
FB1_H
FB3_NL
FB3_H
Type
RMRP
CCN2
STC1
PLCG2
NOVA1
CCL21
CYR61
PTGDS
TNC
APOD
ID1
NR4A1
ID3
IGF2
CD74
FTH1
PCSK1N
IGFBP3
EDNRA
POSTN
CRABP1
TNN
TFAP2A
SFRP2
Gene
Percent Expressed
0
25
50
75
100
H
Type
Neuron_H
Neuron_NL
CLU
CCN1
CCN5
TTR
S100B
NRXN1
SCN7A
DKK3
CAPS
CD9
GSN
RARRES2
GFRA3
CST3
SEMA3B
PDLIM4
PLPP1
Gene
*
**
-0.4
Percent Expressed
0
25
50
75
100
I
IL31
Expression Level
0.12
0.09
0.06
0.03
0.00
CD4Trm
Identity
H
NL
J

**Fig. S6. Differential single cell landscape of non-lesional PN vs healthy skin reveals systemic fibroblast dysfunction**. **a,** Compositional analysis of cell clusters showing differential cell loading coefficients of FB and PCs in NL compared to HC skin. **b,** Barplot displaying composition of cell types in NL and HC skin. Statistical analysis done via Wilcoxon rank sum test with p values shown in adjacent bar graph. **c,** Hierarchical representation of compositional changes in NL and HC skin. **d,** cluster free compositional changes based on subtraction (top) and wilcox testing (bottom). **e,** Expression differences calculated using normalized expression distance between NL and HC skin for all cell clusters. **f,** Gene ontology heatmaps of top 50 pathways upregulated or downregulated in PN L vs HC skin. **g,** Dotplot displaying DEGs for FB1, FB3, and FB7 NL populations displaying a systemic dysregulation compared to HC skin. **h,** Dotplot of DEGs for NL skin neurons compared to HC skin neurons. **i,** Split violin plot displaying increased pruritogenic IL31 expression in CD4Trm in NL skin compared to HCAll dot plot comparisons between H and NL gene expression at a cellular level represent $p<0.05$ using wilcoxon rank sum test. All asterisks represent multiple t tests at a sample level using mean cellular expression, with *$p<0.5$ and ** $p<0.01$.

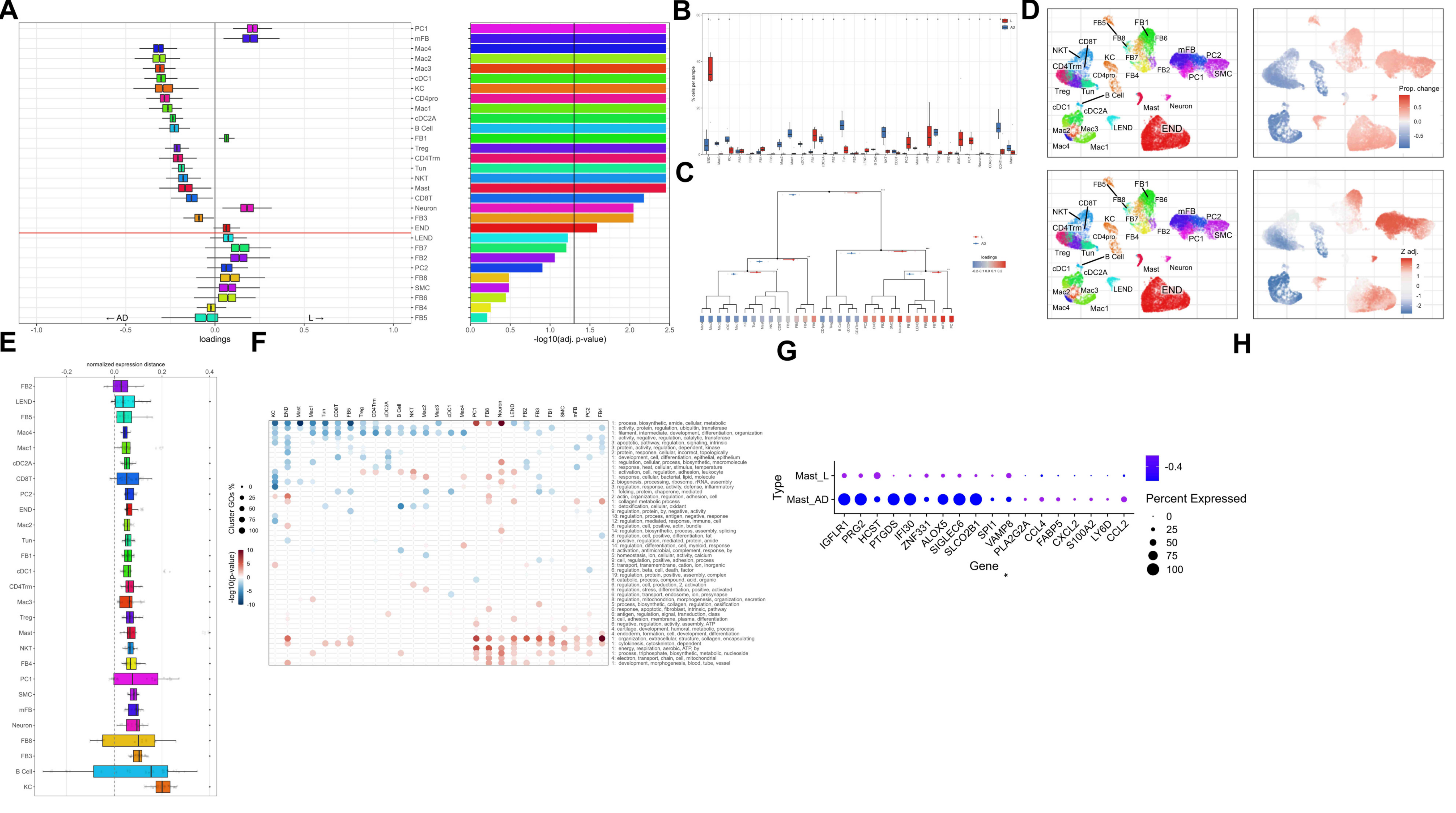
A
B
C
D
E
F
G
H
loadings
← AD
L →
-log10(adj. p-value)
PC1
mFB
Mac4
Mac2
Mac3
cDC1
KC
CD4pro
Mac1
cDC2A
B Cell
FB1
Treg
CD4Trm
Tun
NKT
Mast
CD8T
Neuron
FB3
END
LEND
FB7
FB2
PC2
FB8
SMC
FB6
FB4
FB5
% cells per sample
loadings
Prop. change
Z adj.
normalized expression distance
Cluster GOs %
-log10(p-value)
Type
Mast_L
Mast_AD
Gene
Percent Expressed
IGFLR1
PRG2
HCST
PTGDS
IFI30
ZNF331
ALOX5
SIGLEC6
SLCO2B1
SPI1
VAMP8
PLA2G2A
CCL4
FABP5
CXCL2
S100A2
LY6D
CCL2

**Fig. S7. Differential single cell landscape of lesional PN vs lesional AD reveals neural-fibroblast predominance in PN and adaptive immunity in AD. a,** Compositional analysis of cell clusters showing differential cell loading coefficients of FB in L compared to immune cells in AD skin. **b,** Barplot displaying composition of cell types in L and AD skin. Statistical analysis done via Wilcoxon rank sum test with p values shown in adjacent bar graph. **c,** Hierarchical representation of compositional changes in L and AD skin. **d,** cluster free compositional changes based on subtraction (top) and wilcox testing (bottom). **e,** Expression differences calculated using normalized expression distance between L and AD skin for all cell clusters. **f,** Gene ontology heatmaps of top 50 pathways upregulated or downregulated in PN L vs AD skin. **g,** Dotplot displaying DEGs for Mast cells in L compared to AD. All dot plot comparisons between H and NL gene expression at a cellular level represent $p<0.05$ using wilcoxon rank sum test. All asterisks represent multiple t tests at a sample level using mean cellular expression, with *$p<0.5$ and ** $p<0.01$.

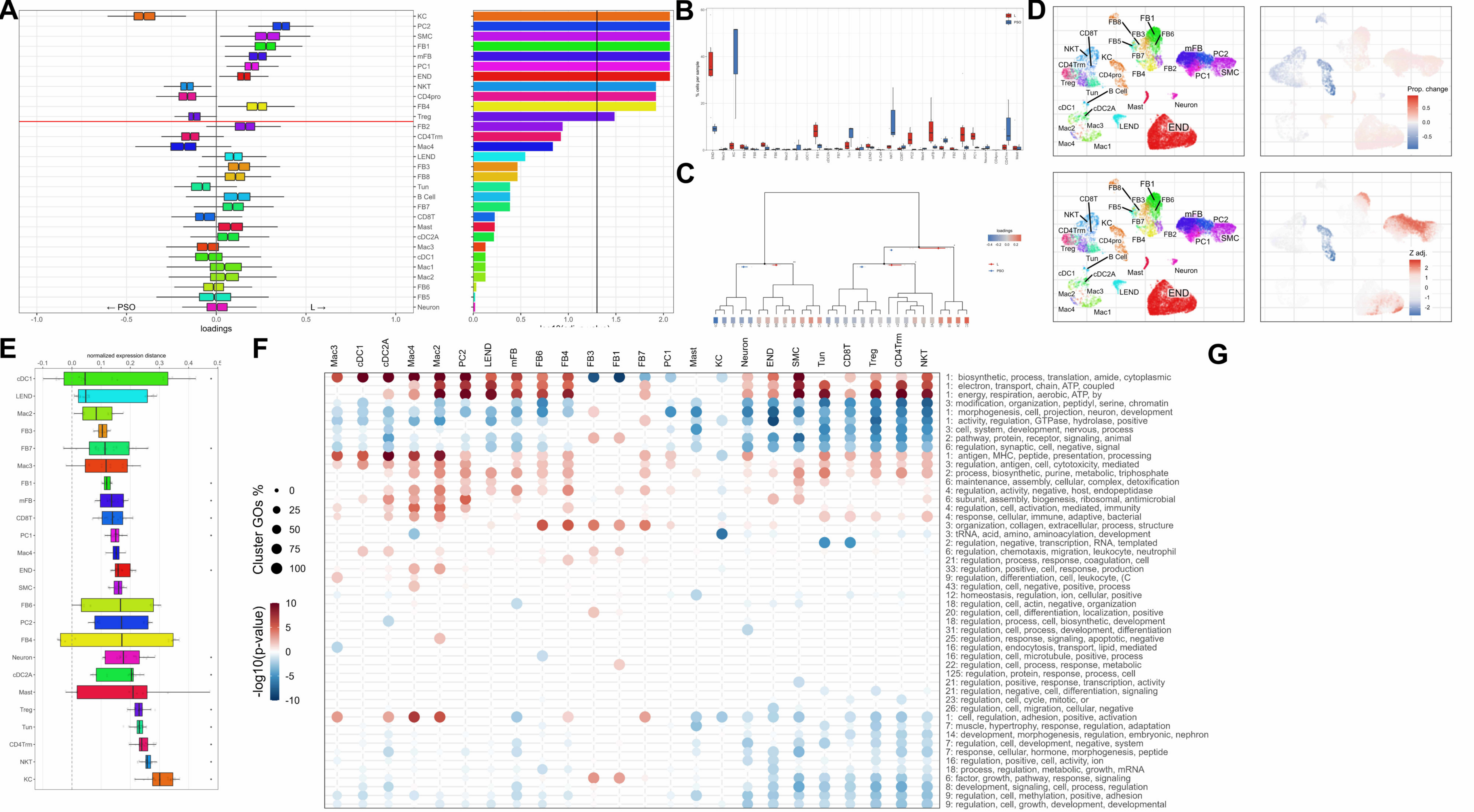

A
KC
PC2
SMC
FB1
mFB
PC1
END
NKT
CD4pro
FB4
Treg
FB2
CD4Trm
Mac4
LEND
FB3
FB8
Tun
B Cell
FB7
CD8T
Mast
cDC2A
Mac3
cDC1
Mac1
Mac2
FB6
FB5
Neuron
← PSO
L →
-1.0
-0.5
0.0
0.5
1.0
loadings
0.0
0.5
1.0
1.5
2.0
B
C
D
FB8
FB1
CD8T
FB3
FB6
FB5
NKT
KC
FB7
mFB
PC2
CD4Trm
CD4pro
FB4
FB2
PC1
SMC
Treg
Tun
B Cell
cDC1
cDC2A
Mast
Neuron
Mac2
Mac3
LEND
END
Mac4
Mac1
Prop. change
0.5
0.0
-0.5
Z adj.
2
1
0
-1
-2
E
normalized expression distance
-0.1
0.0
0.1
0.2
0.3
0.4
0.5
cDC1
LEND
Mac2
FB3
FB7
Mac3
FB1
mFB
CD8T
PC1
Mac4
END
SMC
FB6
PC2
FB4
Neuron
cDC2A
Mast
Treg
Tun
CD4Trm
NKT
KC
F
Mac3
cDC1
cDC2A
Mac4
Mac2
PC2
LEND
mFB
FB6
FB4
FB3
FB1
FB7
PC1
Mast
KC
Neuron
END
SMC
Tun
CD8T
Treg
CD4Trm
NKT
Cluster GOs %
0
25
50
75
100
-log10(p-value)
10
5
0
-5
-10
1: biosynthetic, process, translation, amide, cytoplasmic
1: electron, transport, chain, ATP, coupled
1: energy, respiration, aerobic, ATP, by
3: modification, organization, peptidyl, serine, chromatin
1: morphogenesis, cell, projection, neuron, development
1: activity, regulation, GTPase, hydrolase, positive
3: cell, system, development, nervous, process
2: pathway, protein, receptor, signaling, animal
6: regulation, synaptic, cell, negative, signal
1: antigen, MHC, peptide, presentation, processing
3: regulation, antigen, cell, cytotoxicity, mediated
2: process, biosynthetic, purine, metabolic, triphosphate
6: maintenance, assembly, cellular, complex, detoxification
4: regulation, activity, negative, host, endopeptidase
6: subunit, assembly, biogenesis, ribosomal, antimicrobial
4: regulation, cell, activation, mediated, immunity
4: response, cellular, immune, adaptive, bacterial
3: organization, collagen, extracellular, process, structure
3: tRNA, acid, amino, aminoacylation, development
2: regulation, negative, transcription, RNA, templated
6: regulation, chemotaxis, migration, leukocyte, neutrophil
21: regulation, process, response, coagulation, cell
33: regulation, positive, cell, response, production
9: regulation, differentiation, cell, leukocyte, (C
43: regulation, cell, negative, positive, process
12: homeostasis, regulation, ion, cellular, positive
18: regulation, cell, actin, negative, organization
20: regulation, cell, differentiation, localization, positive
18: regulation, process, cell, biosynthetic, development
31: regulation, cell, process, development, differentiation
25: regulation, response, signaling, apoptotic, negative
16: regulation, endocytosis, transport, lipid, mediated
16: regulation, cell, microtubule, positive, process
22: regulation, cell, process, response, metabolic
125: regulation, protein, response, process, cell
21: regulation, positive, response, transcription, activity
21: regulation, negative, cell, differentiation, signaling
23: regulation, cell, cycle, mitotic, or
26: regulation, cell, migration, cellular, negative
1: cell, regulation, adhesion, positive, activation
7: muscle, hypertrophy, response, regulation, adaptation
14: development, morphogenesis, regulation, embryonic, nephron
7: regulation, cell, development, negative, system
7: response, cellular, hormone, morphogenesis, peptide
16: regulation, positive, cell, activity, ion
18: process, regulation, metabolic, growth, mRNA
6: factor, growth, pathway, response, signaling
8: development, signaling, cell, process, regulation
9: regulation, cell, methylation, positive, adhesion
9: regulation, cell, growth, development, developmental
G

**Fig. S8. Differential single cell landscape of lesional PN vs lesional AD reveals fibroblast predominance in PN and epithelial-immune dysregulation in PSO. a,** Compositional analysis of cell clusters showing differential cell loading coefficients of FB in L compared to KC and T cells in PSO skin. **b,** Barplot displaying composition of cell types in L and PSO skin. Statistical analysis done via Wilcoxon rank sum test with p values shown in adjacent bar graph. **c,** Hierarchical representation of compositional changes in L and PSO skin. **d,** cluster free compositional changes based on subtraction (top) and wilcox testing (bottom). **e,** Expression differences calculated using normalized expression distance between L and PSO skin for all cell clusters. **f,** Gene ontology heatmaps of top 50 pathways upregulated or downregulated in PN L vs PSO skin. **g,** Split violin plots of DEGs in CD4Trm (IL13, IL17A) revealing Th2 shift in PN compared to PSO. Statistical analysis of PSO vs PN L gene expression at a cellular level represent $p<0.05$ using wilcoxon rank sum test.

To view complete supplementary figures including volcano plots visit Mendeley database at:
Patel, Jay (2023), “Single-cell RNA sequencing reveals dysregulated fibroblast subclusters in prurigo nodularis”, Mendeley Data, V1, doi: 10.17632/stmb2m7sb5.1
